# Circadian temperature compensation is intrinsically linked with metabolism and redox signaling

**DOI:** 10.64898/2026.06.23.734026

**Authors:** Yao Xu, Tetsuya Mori, Himanshu Mehra, Alessandro Ustione, Kenya Tanaka, Yuta Kitaguchi, Koichiro Uriu, Shuji Nakanishi, David W. Piston, David Vinyard, Carl Hirschie Johnson

## Abstract

Circadian rhythms are a universal property of organisms, and a defining property of these oscillators is the ability to maintain a precise period at different temperatures. Indeed, the discovery in the 1950s of this property, known as “temperature compensation,” was a watershed event in our understanding of biological timekeeping. Since that time, however, almost no general principles have emerged that uncover the mechanistic basis of how circadian timekeepers defy the ‘rules” of temperature dependence of biochemical reactions. A genetic investigation of circadian temperature compensation revealed that the sensing of metabolism and ensuing modulation of the core mechanism preserves a constant circadian period. Intracellular redox sensing of metabolic rate is intrinsic to this compensation, and this relationship is conserved in bacterial and mammalian cells. These insights explain previously inexplicable observations of the interaction between redox reactions and circadian timekeeping. These results introduce a previously unknown linkage between temperature compensation and metabolism.

## Introduction

Circadian rhythms are biological timekeeping mechanisms that provide a fitness-enhancing internal clock to anticipate of time-of-day changes in the environment.^1–3^ To accomplish this, appropriate phase alignment of these rhythms with the environment is essential; the period of circadian clocks must be homeostatically buffered against changes in the intra-organismal and extra-organismal environments in order to maintain the constant period length that is necessary for accurate phase alignment.^1,4,5^ This includes buffering against changes in temperature within the physiological range, to which biochemical reactions are invariably sensitive, with Q_10_ values typically in the range of 2 to 3. On the other hand, most circadian rhythms have Q_10_ values between 0.9 and 1.1, and a hypothetical model of counterbalancing enzymatic reactions was proposed in 1957 to account for this temperature compensation (TC) property (Extended Data Fig. 1a).^6^ Generally based upon this decades-old hypothesis, the current explanatory mechanisms for TC invoke inter-protein interactions such as phosphorylation^7–10^ or differential gene expression,^10^ but these mechanisms do not adequately explain the full range of circadian compensation phenomena.

Because circadian mechanisms depend upon cellular energy, changes in metabolism can also influence the accuracy of timekeeping. Indeed, changes in metabolism are known to affect circadian free-running period (FRP).^11–17^ Reciprocity between the circadian clockwork and metabolic functions has been documented in organisms at every level of cellular organization.^9–16,18,19^ Moreover, the FRPs of some circadian systems are additionally compensated for altered nutritional conditions.^10^ This “Nutritional Compensation” appears to be regulated by transcriptional, post-transcriptional, and post-translational mechanisms involving metabolism.^10^ Temperature has large effects on metabolic rates, but the possibility that circadian compensation for temperature and metabolic changes might have an underlying common linkage has not been considered. To illuminate the mechanism of temperature compensation (TC), we undertook an unbiased genetic screen in a model system for mutants of the circadian clock that are defective in this property. Our genetic screen identified mutants that revealed a heretofore unappreciated connection between the sensing of metabolic rate and the circadian compensation mechanism for temperature.^17^

## Results

### Genetic screen for temperature compensation mutants reveals sensitivity to metabolism

Our unbiased genetic screen for TC mutants of the circadian clock used the model circadian system *Synechococcus elongatus*.^20–22^ We mutagenized by error-prone PCR the central circadian gene cluster *kaiABC* (Extended Data Fig. 1b)^21^ and initiated a high-throughput screening^22^ of the circadian rhythm of luminescence at 25°C and 30°C to identify clones that were defective in temperature compensation of the free-running period (FRP) in constant light (LL, Fig. 1a & Extended Data Fig. 1c). This strategy identified a wide range of mutant strains whose FRP was dramatically different at 25°C as compared with at 30°C, ranging from Q_10_ values of 0.6 to 1.6 (Extended Data Fig. 1c-e), while the Q_10_ of wild-type (WT) is ∼1.07 (Fig. 1a). To further characterize these temperature-compensation mutants (TC-mutants), we tested their response to dim light intensities (10 μE m^-2^ sec^-1^), a condition which has negligible impact on the FRP of WT. To our surprise, we found that the FRP of many of the TC-mutants was dramatically altered in dim light (Fig. 1d, Extended Data Fig. 2a,c). Because *S. elongatus* is photoautotrophic, reducing the intensity of light substantially slows growth rate and metabolic functions, as does lowering the temperature (Extended Data Fig. 3). We therefore considered the possibility that FRP of the TC-mutants might be more sensitive than WT to metabolic perturbations in general. Indeed, we found that the FRPs of the TC-mutant strains were also more sensitive than WT to the addition of a wide range of metabolic inhibitors including mannitol (a metabolizable sugar alcohol), the glucose analog 3-*O*-methylglucose, the translation inhibitor kasugamycin, and the photosynthesis inhibitor dibromothymoquinone (DBMIB, Fig. 1d, Extended Data Fig. 2). These metabolic conditions slow growth rate of WT cells (Extended Data Fig. 3), but the FRP of WT is practically unaffected (Fig. 1d, Extended Data Fig. 2a), so the intact circadian system in WT is not only compensated for the effects of temperature, but also for metabolic rates.

**Fig. 1.**
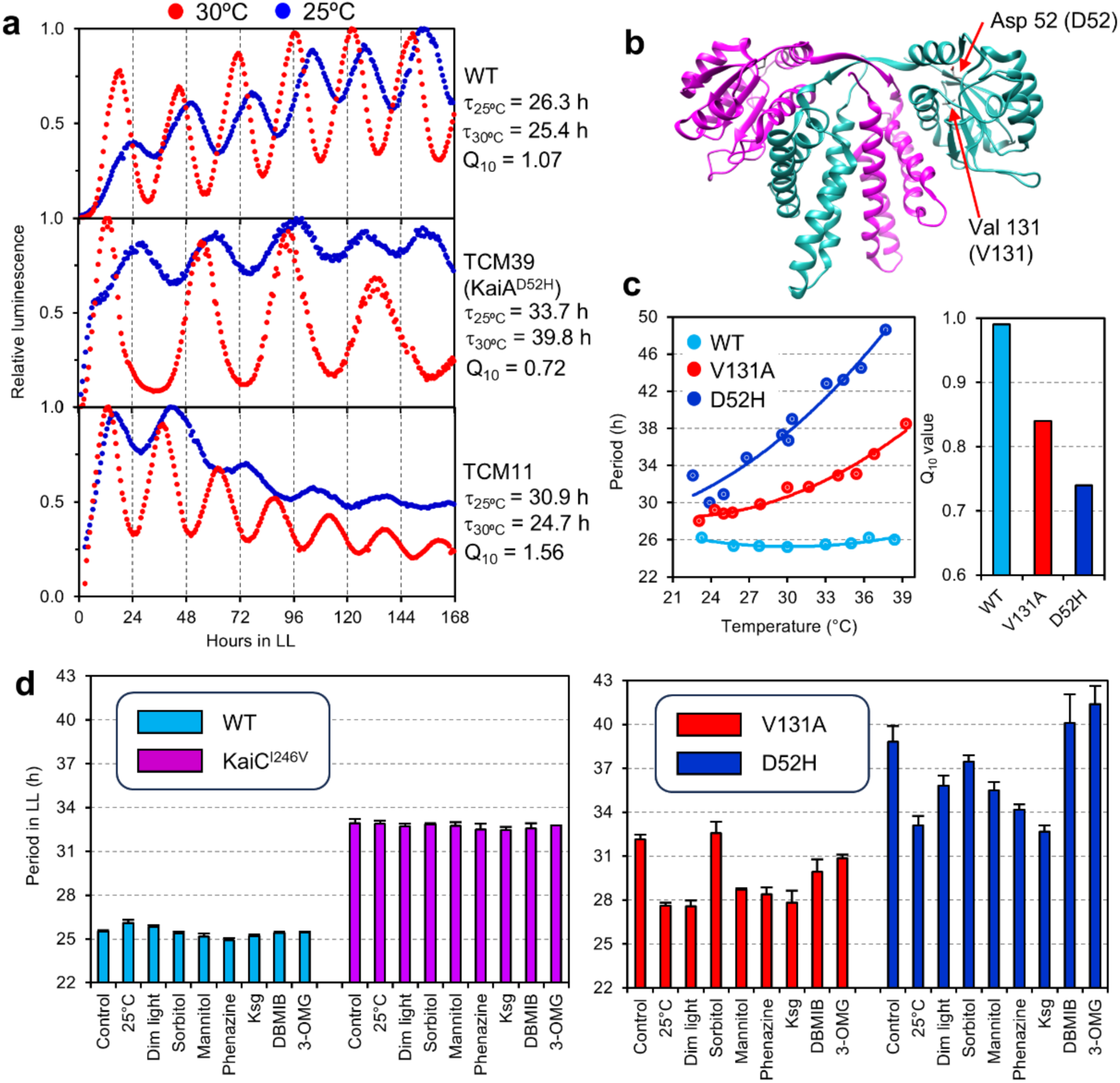
Identification of mutations in core clock gene *kaiA* that are defective in both temperature and metabolic compensation. **a.** Examples of two temperature compensation mutants (TCM11 and TCM39) *vs.* wild type (WT): rhythms with free running period (FRP, τ) and Q_10_ values at 25°C and 30°C, respectively. **b.** Location of two key TC mutations in the KaiA structure. **c.** Temperature dependency of FRP for WT and two KaiA mutants (V131A and D52H) over a broad range of temperatures (left). The Q_10_ values calculated from these data are shown in the right panel. **d.** FRPs of WT, KaiC^I246V^, KaiA^V131A^ and KaiA^D52H^ strains at 30°C in the absence or presence of metabolic inhibition: lower temperature (25°C), dim light (10 μE m^-2^·sec^-1^), 3% sorbitol, 3% mannitol, 20 μM phenazine, 2 μg mL^-1^ kasugamycin (Ksg), 30 μM DBMIB, and 0.2% 3-OMG. The long period mutant KaiC^I246V^ is well compensated for altered temperature or metabolism (left), but the KaiA TC-mutants V131A and D52H are not (right). Error bars are S.D., n = 3.

Among the pool of TC mutants, we identified four point mutations in the *kaiA* clock gene (Fig. 1b, Extended Data Fig. 4) that were particularly interesting because the strains harboring these *kaiA* mutations (i) exhibited the unusual property of “over”-compensation for temperature, i.e., the FRP was *faster* at colder temperatures (Q_10_ < 1.0; Fig. 1c & Extended Data Fig. 4b,c), and (ii) had significantly altered FRPs in response to the metabolic conditions (Fig. 1d; Extended Data Fig. 2a). We focused upon two of these mutants (D52H & V131A, Fig. 1b) because their Q_10_ < 1.0 phenotype and responses to temperature and metabolic conditions were reproducible and robust (Fig. 1c,d). Although these *kaiA* TC-mutant strains had intrinsic FRPs that were longer than that of WT (WT FRP is ∼25 h while the kaiA mutant FRPs were 30-38 h at 30°C), long FRP is not necessarily correlated with impaired compensation for temperature and metabolic rate; e.g., another long FRP strain (KaiC^I246V^, FRP∼33 h) showed WT-like compensation for temperature changes and metabolic inhibition (Fig. 1d, Fig. 2a).

**Fig. 2.**
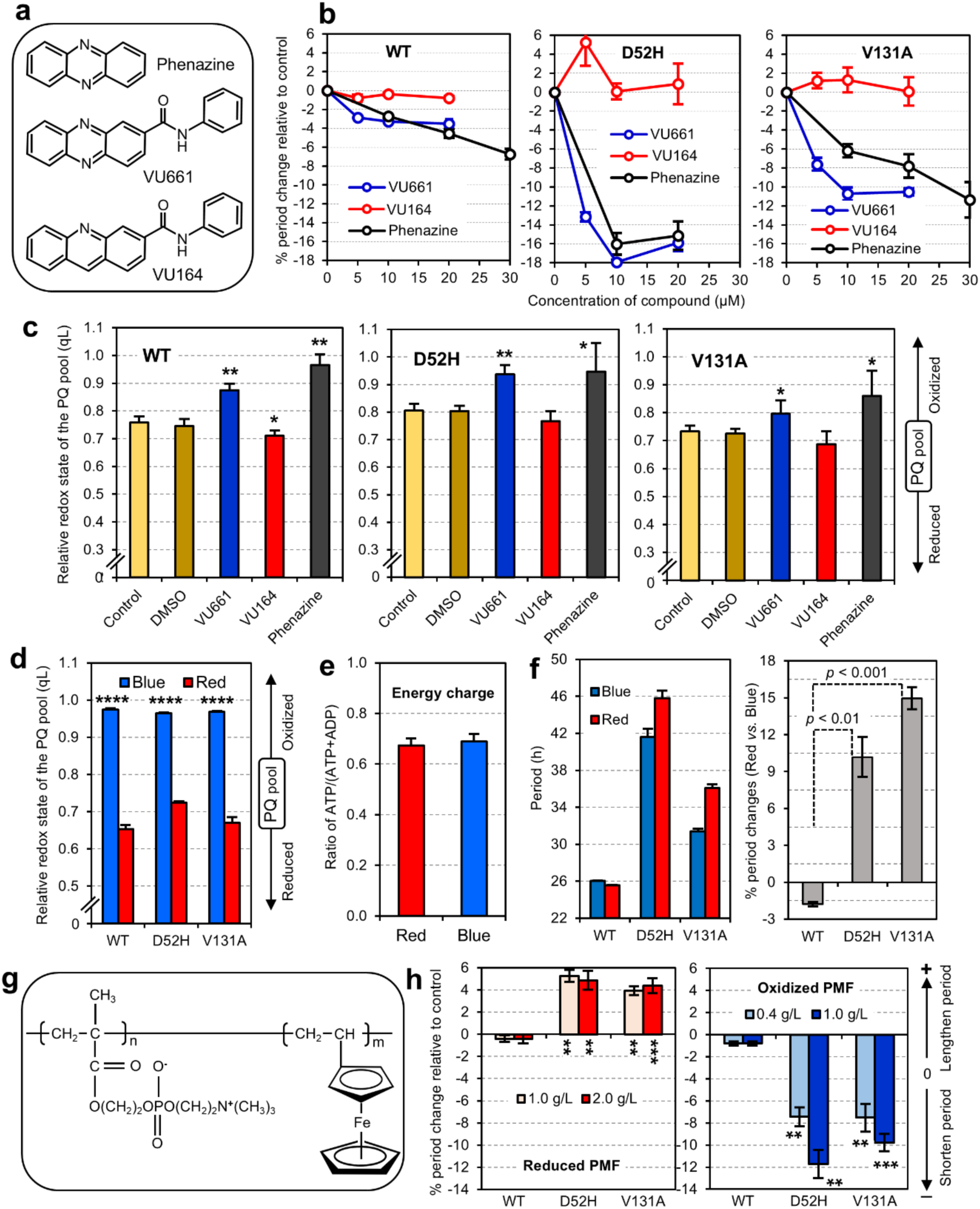
Temperature Compensation *kaiA* Mutants are Sensitive to redox changes *in vivo.* **a.** Chemical structures of phenazine and its derivatives VU661 and VU164. **b.** Electron-shuttling phenazine and VU661 shorten FRP in the TC mutants of KaiA. Differences in FRP among WT, KaiA^D52H^ and KaiA^V131A^ strains under different concentrations of redox drugs are normalized by expressing changes in FRP as “% change relative to control,” where negative values are FRP shortening, and positive values are FRP lengthening (n = 3). **c.** Redox state estimated as the fraction of open PSII reaction centers (q_L_) among WT, KaiA^D52H^ and KaiA^V131A^ strains in the absence (Control and dimethyl sulfoxide [DMSO]) or in the presence of 50 μM VU661, 50 μM VU164, or 30 μM phenazine for 24h. The larger q_L_ values indicate a more oxidized state of the PQ pool, whereas smaller q_L_ values indicate a more reduced state (see Methods) (n = 3-5). **d.** Comparison of redox states (q_L_) under red *vs.* blue light conditions between WT and *kaiA* TC-mutants (n = 3). **e.** Energy charge status [ratio of ATP/(ATP+ADP)] of WT strain under red and blue light conditions (n = 3). **f.** FRP values expressed in hours (left) and % changes (right) for the WT, KaiA^D52H^ and KaiA^V131A^ strains under red *vs*. blue light. *p* values are compared to the WT data (n = 3). **g.** Chemical structure of poly-2-methacryloyloxyethyl phosphorylcholine-co-vinylferrocene (PMF). **h.** Opposite effects of the reduced *vs.* oxidized PMF on FRP of *kaiA* TC-mutants. Reduced PMF lengths FRP of the *kaiA* TC-mutants, whereas oxidized PMF shortens the FRPs (n = 3). Error bars are mean ± S.D., * p<0.05, ** p<0.01, *** p<0.001, ****p<0.0001 by t-tests or Two-way ANOVA.

### Compensation mechanism involves redox sensing

Altered metabolic status is associated with changes of intracellular redox,^18,23,24^ and KaiA alters circadian phenotypes in response to changes of redox.^25,26^ We set out to test if these correlations were related to the phenomena we observed with KaiA TC-mutant strains and temperature compensation. We recently reported that redox-active phenazine analog molecules can affect FRP in mammalian cells by disrupting redox homeostasis.^27^ Phenazine and the phenazine carboxamide VU661 are electron-shuttling compounds and the FRP of the *kaiA* TC-mutants is significantly more sensitive than WT to these compounds; on the other hand, the redox inactive analog of VU661, VU164, has little effect on FRP (Figs. 1a, Fig. 2a,b). To confirm that these treatments affect redox, we estimated overall cellular redox by measuring the redox status of the plastoquinone (PQ) pool that is shared in cyanobacteria by photosynthesis and respiration,^28^ and is measurable by chlorophyll fluorescence (see Methods). We found that phenazine and VU661 oxidize the PQ while VU164 has no effect (Fig. 2c). The redox state can also be altered non-pharmacologically and non-invasively in cyanobacteria by growth in constant red (RR) or constant blue (BB) light.^29^ RR leads to PQ pool reduction while BB leads to PQ pool oxidation (Fig. 2d), while the Energy Charge (ATP/{ATP+ADP}) is nearly the same in red *versus* blue light (Fig. 2e). These effects on PQ redox correlate with FRP: the FRP of WT is practically unaffected by the color of light, whereas the FRP of KaiA^D52H^ and KaiA^V131A^ are consistently longer under the reducing condition of RR as compared with BB (Fig. 2f). These data with phenazines and light color suggest that oxidizing conditions shorten–whereas reducing conditions lengthen–the FRP of the KaiA TC-mutants while WT cells are minimally affected.

This correlation is confirmed by controlling the redox of the cells directly by extracellular electron transfer, a process in which the intracellular electrons of cyanobacteria are exchanged across the cell membrane. We tested the transmembrane biocompatible electron mediator, poly-2-methacryloyloxyethyl phosphorylcholine-co-vinylferrocene (PMF, Fig. 2g),^30^ by providing PMF in a reduced vs. oxidized state in the medium, the intracellular redox of cyanobacterial cells can be reduced vs. oxidized, respectively.^31^ Using this direct method to control redox status, we found that reduced PMF lengthens, while oxidized PMF shortens the FRPs of the *kaiA* TC-mutant strains (Fig. 2h; fluorescence measurements of PQ status during PMF treatment were not possible due to competing PMF fluorescence). On the other hand, the FRP of WT cells is almost unaffected by the redox changes mediated by PMF (Fig. 2h). Taken together, these various treatments that modulate redox by different actions indicate that the FRP of the *kaiA* TC-mutant strains *in vivo* is shortened by intracellular oxidation and lengthened by reduction, whereas the FRP of WT is well compensated against these changes.

### *In vitro* circadian oscillator is also compensated against redox changes

The cyanobacterial circadian system uniquely enables the analysis of an *in vitro* oscillator (IVO) that reconstitutes a core Post-translational Oscillator (PTO) that operates *in vivo*.^20,32,33^ The minimal components necessary to comprise this IVO are KaiA, KaiB, KaiC, and ATP.^32^ The role of KaiA in this oscillatory mechanism is primarily to promote KaiC’s autophosphorylation,^34,35^ and therefore we tested whether this action of KaiA was impaired in the KaiA TC-mutants *in vitro*. A slower rate of KaiA-stimulated KaiC autophosphorylation was characteristic of the KaiA mutants (Fig. 3a, Extended Data Fig. 5). Moreover, the amplitude of the IVO oscillator is dramatically reduced and the IVO period is slightly shortened when KaiA^WT^ in the IVO is replaced with the mutant KaiA proteins (Fig. 3b); these amplitude and period effects in the IVO can be simulated on the basis of the reduced KaiC autophosphorylation activity (Fig. 3c,d; Extended Data Fig. 5d,e). Both KaiA and CikA are sensitive to redox,^26,36^ and the KaiA mutants are more strongly affected than KaiA^WT^ by oxidation from the quinone analog Q_o_ both in terms of KaiC autophosphorylation (Fig. 3e) and damping of the IVO rhythm (Fig. 3f). Therefore, consistent with the studies of intact cells (Figs. 1 & 2), the core biochemical IVO is sensitized to perturbations of redox by the TC-mutations in KaiA. *In vivo*, this IVO functions as a PTO that triggers a global Transcription/Translation Feedback Loop (TTFL) through the action of CikA, SasA, and RpaA (Fig. 3c) to enable the rhythmic gene expression monitored by the luminescent *luxAB* reporter (Fig. 1a).^20,33^ Our integrated PTO/TTFL model can simulate the longer periods of the KaiA mutant strains *in vivo* at 30°C (Fig. 1c,d) on the basis of the less active KaiA (Fig. 3a) that leads to a reduced amplitude of the IVO/PTO (Fig. 3b). Moreover, oxidation suppresses CikA levels *in vivo*^36^ (Extended Data Figs. 6 & 7), allowing enhanced phosphorylation of the global transcriptional factor RpaA and therefore a stimulation of the *kaiC* transcription feedback loop (Fig. 3c). This energized feedback loop shortens the FRP of the integrated PTO+TTFL oscillation (Fig. 3d), thereby counterbalancing the *in vivo* period-lengthening effects of the KaiA mutants (Figs. 1d, 2, 3d, Extended Data Fig. 7). The final outcome *in vivo* is that (i) the less active KaiA mutants cause a longer period at 30°C, and (ii) redox-oxidizing conditions suppress CikA activity and dampen the IVO/PTO composed of the sensitized mutant KaiA, thereby shortening the observed FRP.

**Fig. 3.**
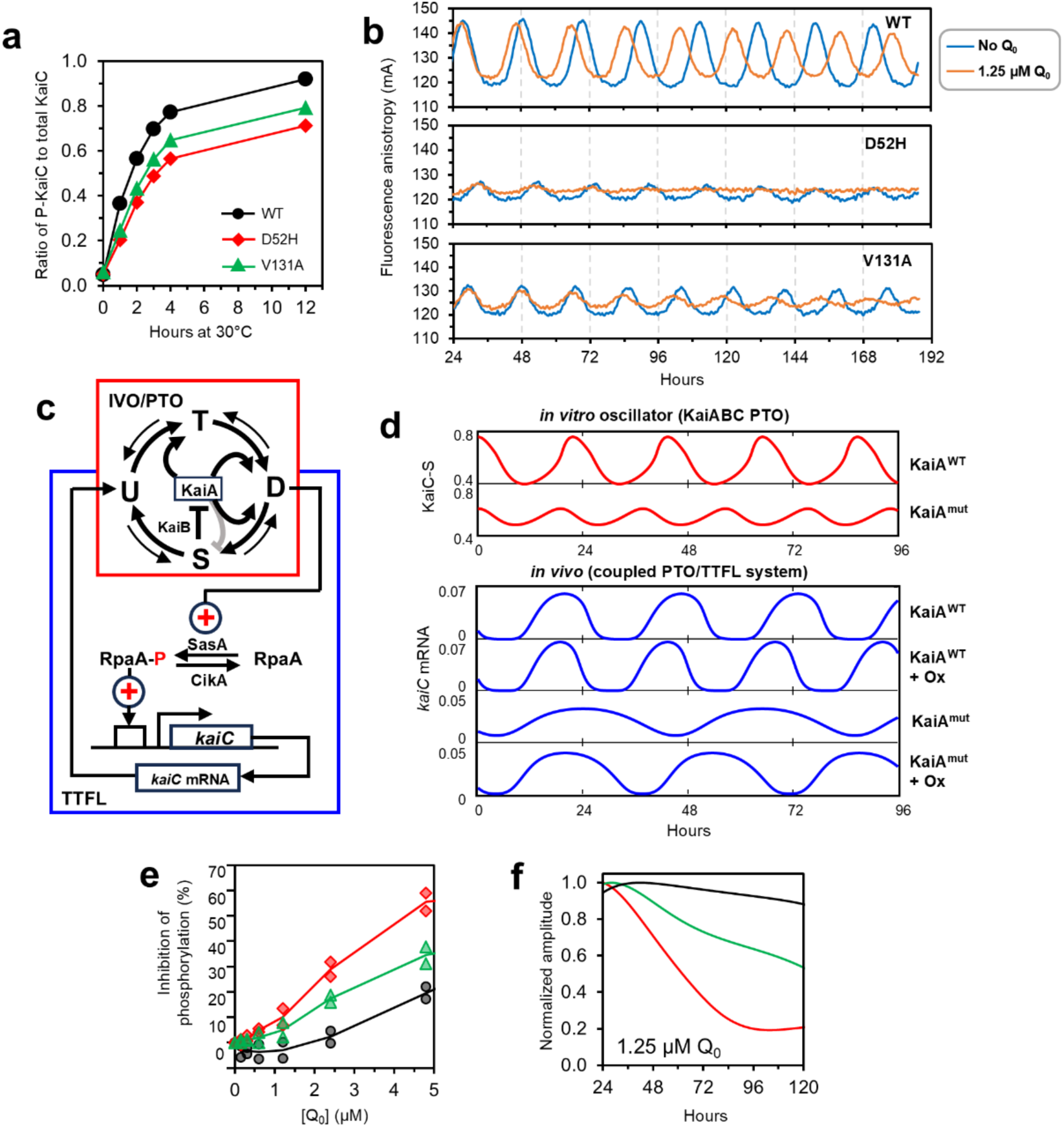
Temperature compensation mutants of KaiA are sensitive to redox changes *in vitro.* **a.** KaiC autokinase activity stimulated by KaiA is reduced by TC mutations. Dephosphorylated KaiC^WT^ was incubated in the presence of the KaiA variants at 30°C. KaiC phosphorylation was assessed by SDS-PAGE (see Extended Data Fig. 5f for gels). **b.** Oscillations of KaiABC complex association/dissociation reconstituted *in vitro* and monitored by fluorescence anisotropy.^33^ Oscillations are shown in the absence or presence of coenzyme Q_0_ (oxidized form, 1.25 μM in this figure, see Extended Data Fig. 5a for other concentrations) (n = 2). **c.** Coupled PTO/TTFL model to simulate the *in vivo* circadian system in cyanobacteria. Post-Translational Oscillator (PTO, red square; phosphorylation states of KaiC: U = 431S/432T, T = 431S/432Tp, D (aka ST) = 431Sp/432Tp, and S=431Sp/T432) and Transcriptional-Translational Feedback Loop (TTFL, blue rectangle) are coupled. The PTO drives the TTFL through a two-component regulatory system consisting of SasA, CikA and RpaA, and there is feedback of newly synthesized KaiC upon the PTO.^64^. **d.** Mathematical simulation of effects of KaiA mutations and redox change on the KaiABC PTO (*in vitro*) and the PTO-TTFL coupled system (*in vivo*). Upper two panels (red traces): rhythms of KaiC phosphorylation state with the wild-type (KaiA^WT^) and variant (KaiA^mut^) KaiAs were simulated by the PTO-only model; ordinate, the abundance of the S (431Sp/T432) form of KaiC. Lower four panels (blue traces): the circadian system *in vivo* was simulated with a coupled PTO-TTFL model, where time-dependent changes of *kaiC* mRNA levels are tracked under standard conditions *versus* oxidized (Ox) conditions (see Extended Data Fig. 7 for details of the simulations). **e.** TC-mutants of KaiA are more sensitive than KaiA^WT^ to oxidation *in vitro*. Purified KaiC and the wild-type or mutant KaiA proteins were incubated at 30°C with various concentrations of coenzyme Q_0_ (0-5 μM). The lines connect the average of two independent experiments. Representative gel images and unnormalized data (including a larger range of Q_0_ concentrations) are shown in Extended Data Fig. 5d,e. **f.** Damping of the amplitude of the *in vitro* oscillations reconstituted with Kai^WT^, KaiA^D52H^, or KaiA^V131A^ in the presence of 1.25 μM Q_0_. The instantaneous amplitudes over time are normalized by the maximum amplitude value for each of the reactions.

### Redox/metabolism compensation mechanism is conserved between bacterial and mammalian cells

Temperature compensation is a defining property of circadian clocks even in cells from endothermic animals.^5,37,38^ The redox-active phenazine carboxamide compound VU661 can also manipulate intracellular redox in mammalian cells at 37°C,^27^ and we extended the impact of VU661 upon intracellular redox in cultured human U2OS cells to both 33°C and 38°C using a noninvasive *ex vivo* measurement of NAD/NADH and FAD/FADH_2_ ratios in cells by Optical Redox Ratio (ORR).^24,27,39^ As shown in Fig. 4a,b, VU661 oxidizes the intracellular redox of human U2OS cells at both 33°C and 38°C as compared with VU164 (VU164 is incapable of serving as an electron shuttle) by decreasing NADH and increasing FAD.^27^ This effect was observed in multiple compartments of the U2OS cells, as microscopic comparisons of ORR across whole cell, mitochondrial, and cytosolic/nuclear compartments showed comparable increases in ORR that result from VU661 treatment (Fig. 4c). Furthermore, comparison of ORR between untreated cells at 33°C versus 38°C indicates significantly increased ORR at 38°C in both the Cell (p = 0.0004) and Cyt/Nuc (p = 0.0002) compartments, confirming that temperature increase is itself oxidizing to intracellular redox status. The disruption of intracellular redox by treatment with VU661 was also observed in a different species of mammalian cells in which the FRP is *over*-compensated for temperature (Rat-1 cells, Q_10_ ∼ 0.8, Extended Data Fig. 8). Moreover, the impact on ORR was equivalent in both cell types at 33°C and 38°C (Fig. 4, Extended Data Fig. 8), so VU661 appears to affect intracellular redox similarly at both temperatures.

**Figure 4.**
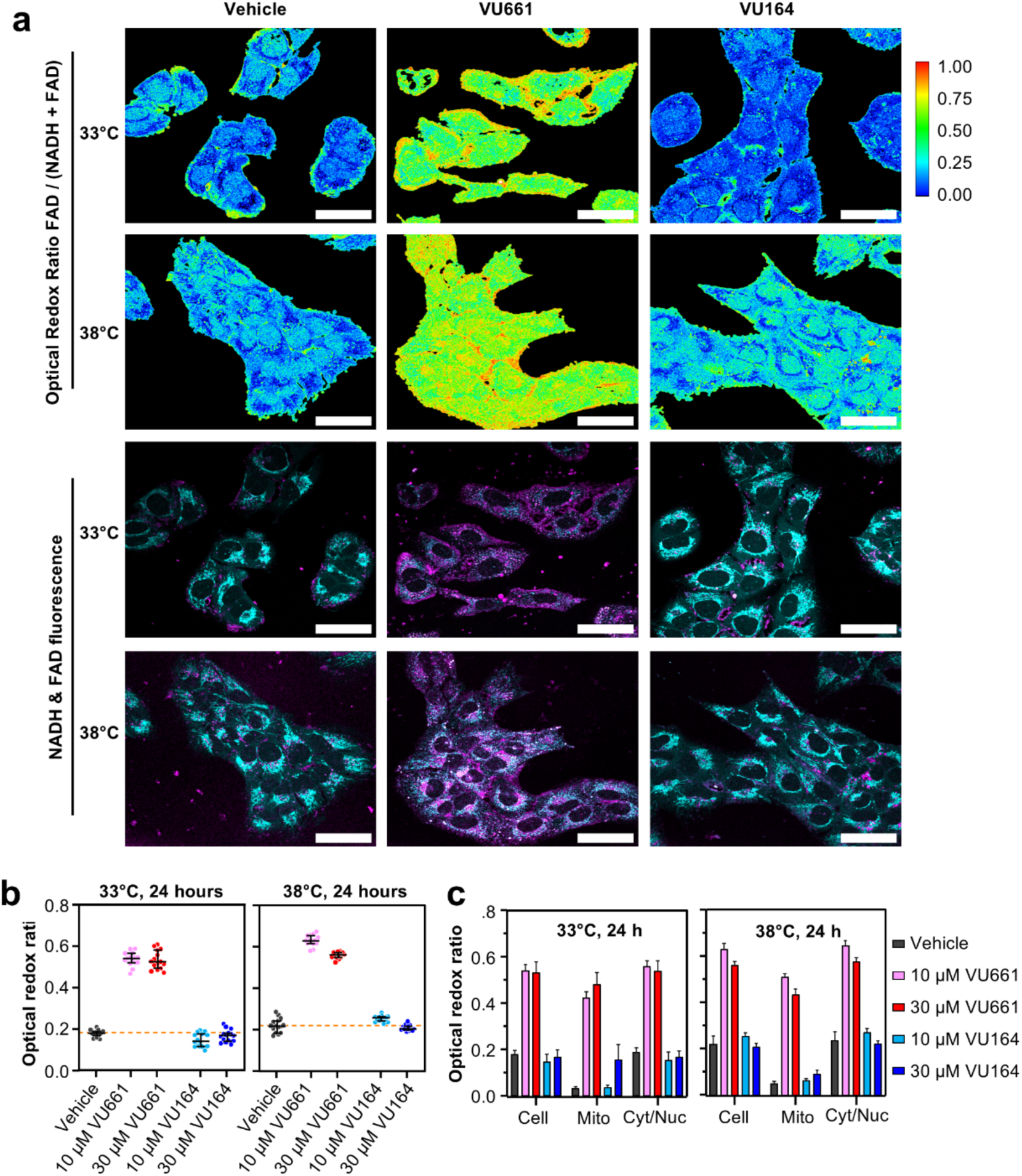
Redox-active phenazine VU661 oxidizes redox in cytosol, mitochondria, and nuclei of mammalian cells at 33°C and 38°C. **a.** Imaging of cellular redox of human U2OS cells following VU661/VU164 treatments at 33°C and 38°C. Images of the Optical Redox Ratio (ORR, top rows), and the corresponding overlays of NADH (cyan) and FAD (magenta) images (bottom rows) after 24 h. One image is displayed for each treatment (vehicle, VU661 10 µM, and VU164 10 µM), and for each temperature (33°C and 38°C). Scale bar 50 µm. **b.** ORR values for all the analyzed images after the 24 h treatment with two concentrations of VU161 or VU164 at 33°C and 38°C, respectively. Each dot in the graph is the value extracted from one image containing 10 – 40 cells, n = 15 for each group. The black bars indicate the median value of each group, and the error bar represent the interquartile range of each group. The dashed line indicates the median value for the control group (vehicle). The NADH and FAD data from which the ORR values in this figure are derived appear in Extended Data Fig. 8d. **c.** ORR data subdivided by cellular compartments: across the whole cell (cell), in the mitochondria (mitochondria), in the non-mitochondria regions (cytoplasm/nucleus). n = 15 for each group, median and interquartile range are displayed.

The application of VU661 reveals that a linkage between redox signaling and circadian temperature compensation is an evolutionarily conserved phenomenon between bacterial and mammalian cells (Fig. 5, Extended Data Fig. 8). In U2OS cells (untreated Q_10_ ∼ 1.1), the temperature compensation properties are significantly impaired by the redox-active VU661 (Q_10_ increases to ∼1.5) but not by its inactive analog VU164 (Fig. 5a,b). In the over-compensated Rat-1 cells treated with VU661, the Q_10_ for circadian FRP also increases from ∼0.79 to ∼0.93 (Extended Data Fig. 8a,b). Similarly, in cyanobacteria, the disruption of temperature compensation by manipulating redox signaling with VU661 is not only observed in the sensitized KaiA-mutant strains (Fig. 2a-c)––a sufficiently strong redox manipulation impairs temperature compensation even in the well-compensated WT strain, as observed by treatment with the redox-active VU661 over a gradient of temperatures in the physiological range (Fig. 5c,d). Therefore, the sensitivity of circadian temperature compensation to redox manipulation is a conserved property of the clock mechanisms of both cyanobacterial and mammalian cells.

**Fig. 5.**
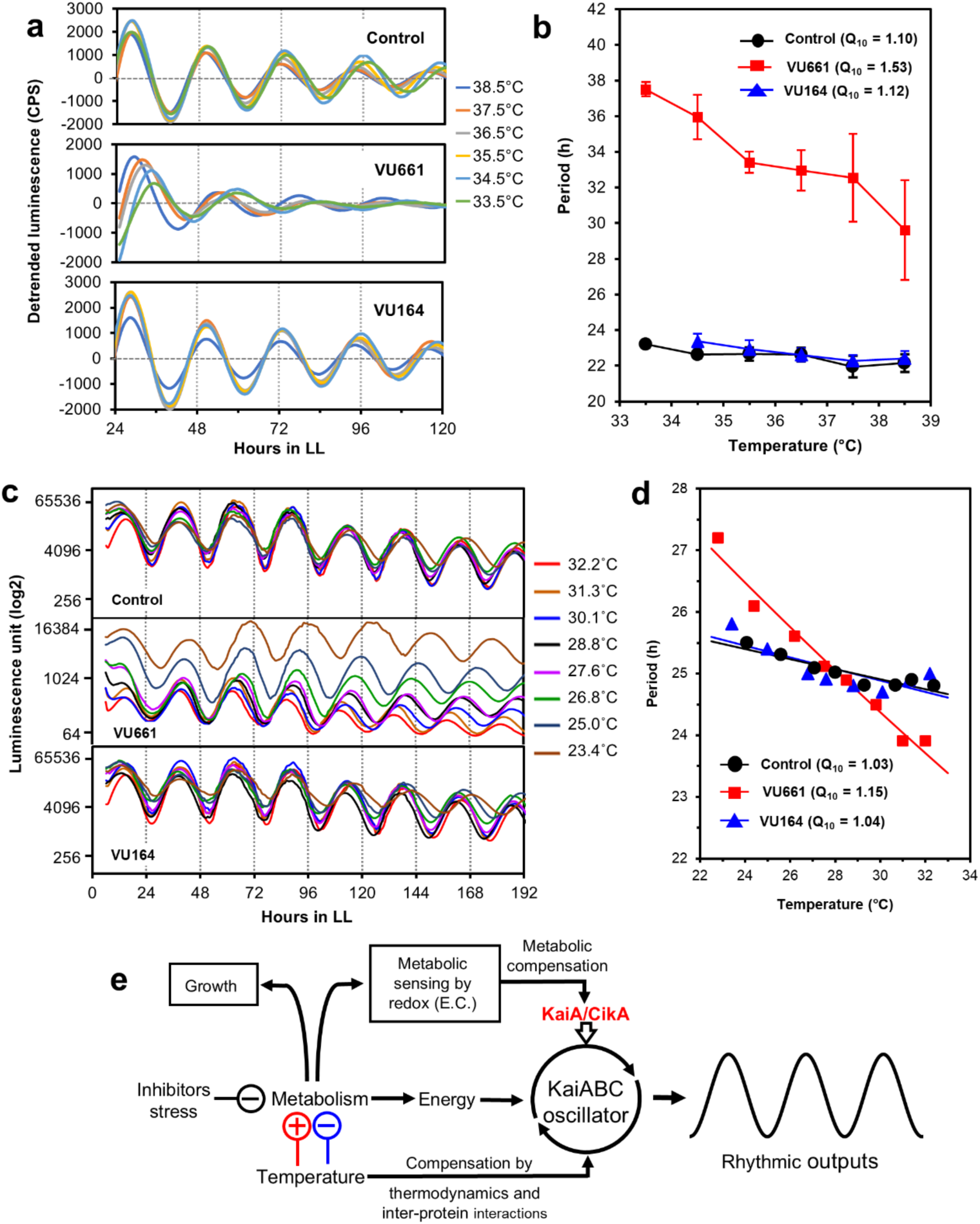
Altering redox status of cyanobacterial and mammalian cells impairs temperature compensation. **a.** Temperature compensation is impaired by the electron-shuttling carboxamide VU661 in human U2OS cells over a temperature gradient of 33.5°C to 38.5°C. U2OS Bmal1-Luc cells were exposed to 30 μM of VU661 or VU164 (+ 0.3% DMSO), while control cells were treated with 0.3% DMSO only. **b.** Periods and Q_10_ values for multiple independent experiments such as in panel b (n = 4). Data are presented as mean ± S.D. for n = 4. **c.** Effect of VU661 and VU164 on free-running circadian rhythms of wild-type cyanobacterial cells *in vivo* over a temperature gradient of ∼ 23°C to ∼ 33°C. **d.** FRP and Q_10_ values of free-running cyanobacterial cells in LL with or without 30 μM of the active form VU661 or an inactive form VU164. Control group contained the same concentration of the vehicle DMSO as that present in the VU661/VU164 treatments. **e.** Model for coupled mechanisms for temperature compensation. Two components of circadian temperature compensation are (i) atomic (thermodynamic properties of each core clock component),^8,44,65^ and (ii) counter-balancing inter-protein interactions.^6^ However, because temperature also affects metabolic production of energy that fuels the circadian clockwork, a third component involves the sensing of metabolism via redox status (and potentially other contributing factors such as Energy Charge {E.C.}) and this modulating information is fed into the core clockwork in cyanobacteria via the redox responsiveness of KaiA and CikA. For a different perspective on counterbalancing compensation and rate-limiting reactions based on redox sensing, see Extended Data Fig. 9.

## Discussion

Why did our genetic screen for TC mutants identify mutations in a compensation mechanism for changes in metabolism? The answer is simple but not obvious. Because it is exquisitely sensitive to cellular temperature, metabolism is a meter of temperature. We find that by enlisting the larger metabolic network, the circadian mechanism circumvents the constraints of temperature-dependent biochemistry, thereby ensuring precise timekeeping. A consequence of this compensation strategy is that unusual perturbations of metabolism can be misinterpreted as temperature changes, so the circadian period can inadvertently become sensitive to such disturbances.^10–14^ While crosstalk between circadian rhythms and metabolism is well appreciated,^9–15^ the intrinsic linkage of metabolism and redox sensing with the circadian temperature compensation mechanism is a new insight.

For the circadian clock to be an accurate timekeeper, it must be buffered against the effects of temperature on the core clock proteins at both atomic (thermodynamic) and inter-protein levels, but it also needs to be compensated against other changes that can occur in the intracellular *milieu* that affect the oscillator. In particular, all known circadian mechanisms require ATP for both energy and as a substrate for phosphorylation reactions. Therefore, circadian oscillators need to be buffered against metabolic changes that might affect ATP availability & concentration. For example, a clock cannot be accurate if it slows down when energy supplies are low. Our results indicate that metabolism is sensed via redox status (and potentially other contributing factors such as Energy Charge) and this modulating information is fed into the core clockwork in cyanobacteria by the redox sensitivity of KaiA and CikA to ultimately conserve FRP (Fig. 5e and Extended Data Fig. 9, model predictions in Fig. 3 and Extended Data Fig. 7). Temperature within the physiological range can influence the intracellular redox status via metabolic rate (thereby altering NAD and NADP redox status, Extended Data Fig. 8c,d) and by the production of reactive oxygen species (ROS). Redox has been previously implicated in circadian properties generally,^11–14,40–42^ but a redox role for temperature compensation is novel. The redox sensors in cyanobacteria include KaiA and CikA; the KaiA TC mutants identified here are hyper-sensitive to redox changes so that the counterbalancing mechanism is inaccurate. Consequently, in cells expressing the mutant KaiA proteins, redox imbalances impair the TC mechanism so that circadian period is no longer conserved across a physiological range of temperatures (Figs. 1, 3, and Extended Data Fig. 9).

In addition to this feedback from metabolism, the other components of circadian temperature compensation rely upon the atomic/thermodynamic properties of the core clock components and counter-balancing inter-protein interactions as proposed 65 years ago (Extended Data Figs. 1a & 9, Fig. 5e).^5–8,14,43^ These three components of temperature compensation – atomic/intra-molecular, inter-molecular, and metabolic – are intrinsically coupled for maximum efficiency. The original hypothesis of counterbalancing INTER-molecular interactions,^6^ especially those involving phosphorylation, has been pursued avidly and several examples have been characterized.^6–8,14,43^ Thermodynamic compensation of individual clock component properties at the atomic level is not as well understood, but there are at least two clock proteins whose activities are temperature compensated in the absence of any inter-protein interaction or metabolic compensation, KaiC and Casein Kinase I.^8,41,43,44^ Metabolic compensation depends upon sensing metabolic status and appropriately modulating the clockwork mechanism to conserve FRP; it is this part of the temperature compensation mechanism that is defective in our KaiA TC mutants such that the circa-24 h FRP is no longer conserved in the face of metabolic *or* temperature perturbations. Previous studies have appreciated that circadian clocks in many organisms are subject to “nutritional compensation,”^10^ but the studies reported herein show that nutritional compensation (what we are calling “metabolic compensation”^17^ is actually a component of the larger homeostatic property that is intrinsically linked with temperature compensation.^17,19^ Moreover, the fact that metabolic compensation of circadian FRP has been observed in a wide variety of organisms^10^ means that this property has been evolutionarily conserved, as has the role of redox sensing (Fig. 5e and Extended Data Fig. 9). A metabolic compensating mechanism based exclusively on sensing ATP levels might seem to be the most direct for the purposes of conserving FRP, however the indirect redox sensor may be a better predictor of impending changes in energy charge. Indeed, our data only show that redox is important, not that it is the exclusive parameter; in fact, our KaiA TC mutants also exhibit some sensitivity to the ATP:ADP ratio (Extended Data Fig. 5f-h).

The significance of these findings is that in order for a circadian clock to be an accurate timekeeper under all intra- and extra-organismal conditions, it must be compensated for the changes of relevant metabolic functions that temperature provokes in addition to its thermodynamic consequences. One consequence of this linkage is that observations of metabolic alterations to circadian period (e.g., by high-fat diet or other nutritional perturbations^11–14^) may be due to interference with the redox sensing component of the temperature compensation mechanism. Our data show that these mechanisms share molecular components in cyanobacteria because mutants in KaiA affect both temperature and metabolic compensation. Another key implication of our study is that it helps to explain contentious observations of circadian redox cycling in mammalian cells and its possible role in licensing circadian-regulated transcription.^45,46^ Circadian redox cycling also occurs in cyanobacteria,^30^ but it is clearly not a component of the core KaiABC oscillator system that drives transcription in these cells^32,33^ (and Fig. 3b here). Many cells undergo daily cycles of metabolism, and these presumably lead to these daily cycles of redox status. We posit that the redox cycling seen in both mammalian and cyanobacterial cells is a manifestation of a compensation loop in action that is keeping track of intracellular metabolism and fine-tuning the clockwork so that FRP is conserved.

## Methods

### Culture conditions and growth rate measurements of cyanobacteria

Cyanobacterial cells of *Synechococcus elongatus* PCC 7942 was grown in modified BG-11 medium^47^ supplemented with appropriate antibiotics (spectinomycin, 20 µg mL^-1^; kanamycin, 10 µg mL^-1^; 5 µg mL^-1^; chloramphenicol, 7.5 µg mL^-1^) under standard conditions of 30°C and 35 ∼ 50 µE m^-2^ s^-1^ (cool-white fluorescence illumination). For red vs. blue light comparison, the light intensity was 35 µE m^-2^ s^-1^ with fixtures of light-emitting diodes (LEDs; 630 nm for red, 460 nm for blue). To evaluate the growth response of the wild type (WT) strain to different metabolic inhibitions, initial cultures were grown in liquid BG-11 medium under standard conditions in a shaking water bath at 100 rpm with air bubbling. When cell densities reached OD_750_ ∼ 0.7, cultures were diluted to OD_750_ ∼ 0.005 or 0.01. The growth rate thereafter was determined by absorbance at 750 nm (OD_750_) daily from cultures in 125-mL flasks containing 70 mL of BG-11 medium in a temperature-controlled shaking water bath at 150 rpm at different temperatures (20°C, 25°C, or 30°C) and under different light intensities (10, 50, or 190 µE m^-2^ s^-1^) in LL or under standard growth conditions in the absence or presence of different concentrations of metabolic treatments, including sorbitol, mannitol, phenazine, kasugamycin, dibromothymoquinone (DBMIB), 3-*O*-methylglucose (3-OMG), or oxidized quinone (Q_0_). Three replicates were performed for each assay.

### Mutagenesis of the *kaiABC* cluster for screening of temperature compensation mutants

A luminescence reporter strain was generated by integrating a *kaiBC*p::luxAB expression cassette into the neutral site I (NS I) of *S. elongatus* PCC 7942 with a chloramphenicol resistance marker in which the expression of the *Vibrio harveyi* luciferase genes *luxAB* are driven by the promoter of the *kaiBC* genes (*kaiBC*p). A *kaiABC*-null strain was created by replacement of the *kaiABC* cluster DNA region with a kanamycin resistance gene. The plasmid pCKaiABC harbors a 3.68 kb of *kaiABC* cluster with 528 bp of *kaiA* 5’-flanking and 288 bp of *kaiC* 3’-flanking regions, in which a 2.02 kb of fragment for spectinomycin resistance marker is inserted to the end of the *kaiC* coding region.^21^ Random mutations were created in the *kaiABC* gene cluster through error-prone polymerase chain reaction (PCR)^48^ with GeneMorph II EZClone Domain Mutagenesis kit (Agilent Technologies, Inc., Santa Clara, CA). Briefly, a population of 2.9 kb mutant megaprimers harboring *kaiABC* cluster DNA were synthesized using Mutazyme II PCR reaction by control mutation rate to about 5 mutations per kb with different initial quantities of wild type *kaiABC* target and amplifying cycles. After purification, the mutated *kaiABC* megaprimers were denatured and annealed to the pCKaiABC plasmid and an EZClone reaction was extended with high fidelity. Plasmids with various mutations in the targeted *kaiABC* cluster were generated in ultracompetent *E. coli* cells, and an efficient *kaiABC* domain mutagenesis library was developed by introducing the mutated *kaiABC* populations with a spectinomycin resistance marker into the endogenous *kai* cluster locus of the *kaiABC*-null strain with replacement of the kanamycin resistance gene, which enables a high-throughput screening in a single experiment for a very broad spectrum of mutations that are specifically targeted to the *kaiABC* cluster (Extended Data Fig. 1b). To use this high-throughput screen for mutations of KaiA, KaiB, and/or KaiC that alter temperature compensation, single colonies generated from the mutagenized *kaiABC* library were first screened for altered FRPs of the luminescence rhythms at lower temperatures (23°C or 25°C) using a turntable “Kondotron” apparatus.^49,50^ The mutant cell lines with different FRPs relative to WT at these lower temperatures were subsequently examined for the FRP phenotype at the higher temperature of 30°C. Based on the FRPs exhibited at these two temperatures (23/25°C vs. 30°C), mutant strains with altered Q_10_ relative to that of WT (Q_10_ = ∼1.07 at 25°C *vs.* 30°C) were saved as potential TC-mutant candidates. After multiple cycles of screening and rescreening at 25°C *vs.* 30°C temperatures, the entire *kaiABC* cluster DNAs from the putative TC-mutant strains were fully sequenced to identify all mutations. These potential TC mutants with single mutation and/or multiple mutations in *kaiABC* region were individually regenerated by site-directed mutagenesis^51^ and introduced as single point mutations to the original *kaiABC* locus of the *kaiABC*-null strain followed by sequence reconfirmation. The regenerated TC-mutants (TCMs) with specific single mutations in *kaiA*, *kaiB*, or *kaiC* were further charactered, evaluated, and compared to the original strains and WT for their temperature compensation properties at different temperatures.

### Preparation of PMF agar medium in different redox states

The redox-active phospholipid polymer, poly-2-methacryloyloxyethyl phosphorylcholine-co-vinylferrocene (PMF, Fig. 2g), was synthesized by free-radical copolymerization of vinylferrocene and 2-methacryloyloxyethyl phosphorylcholine, as described previously.^52^ Because the iron atom in vinylferrocene is in the reduced Fe^2+^ state, the resulting PMF is initially in the reduced form. To obtain the oxidized form of PMF, FeCl_3_ was added to a 40 g L^-1^ aqueous solution of reduced PMF to a final concentration of 0.2 M. The solution was then subjected to dialysis for 9 days using a cellulose membrane with a molecular weight cutoff (MWCO) of 3,500 to remove FeCl_3_. PMF-containing agar plates were prepared by mixing BG-11 agar medium with either the reduced or oxidized PMF solution.

### Measurement of bioluminescence rhythms

For measurement of bioluminescence rhythms of cyanobacterial cells, the *in vivo* luminescence rhythms from single colonies or tooth-picked colonies grown on 1.5% agar BG-11 on plates or vials were continuously monitored at different temperatures or light intensities with or without metabolic treatments using a custom-designed turntable/CCD camera “Kondotron” apparatus or a temperature-controlled cart-moving “Taylortron” apparatus with a photomultiplier tube as described previously.^49,50^ Cells within the populations/colonies were synchronized before measurement of luminescence rhythms by exposure to 1 or 2 cycles of 12h light/12h dark. The free running period (FRP) of luminescence rhythms was analyzed with ChronoAnalysis II, version 10.1 (courtesy of Dr. Till Roenneberg), and the Q_10_ value for evaluation of temperature compensation was calculated as described below.

For measurement of bioluminescence rhythms in mammalian cells, U2OS or Rat-1 cells harboring a Bmal1::Luc reporter^53,54^ were inoculated into 20-mL glass scintillation vials (poly-D-lysine coated bottom surface) containing 2 mL of Dulbecco’s Modified Eagle Medium (DMEM, Gibco 11995-065) with 10% fetal bovine serum (FBS, Gibco A5669701) and 1x Antibiotic-Antimycotic (Gibco 15240062) and grown at 37°C under 5% CO_2_ for a few days. When cultures reached ∼90% confluency, the cells were treated with 100 nM dexamethasone (Dex) in the same growth medium for 2 hours. After the Dex treatment, the medium was replaced with 2 mL of a bioluminescence recording medium containing 10% FBS, 1x Antibiotic-Antimycotic, and 100 μM luciferin.^55^ Compounds VU661 and VU164^27^ were added at 30 μM for U2OS cells or 20 μM for Rat-1 cells to the recording medium. Bioluminescence was monitored with a “Taylortron” custom-made photon-counting apparatus that allows a gradient of temperatures to be presented to a linear array of samples in a single experiment.^49,56^ The assay temperatures were binned in 1°C intervals (*e.g.,* 36.0-36.9°C = 36.5°C) for analysis and presentation.

### Assay of KaiA-stimulated KaiC autokinase activity

The Kai proteins were expressed as GST-fusions in *E. coli* and purified as previously described.^35^ Prior to the assay, 6-9 μM purified KaiC protein was dephosphorylated by incubation at 30°C in assay buffer (20 mM Tris-HCl, pH 8.0, 150 mM NaCl, 5 mM MgCl_2_) containing 1 mM ATP for 36-48 hours. After the incubation, ADP was removed from the resulting solution of dephosphorylated KaiC by application to a desalting spin-column (Zeba Spin Desalting Columns 7K MWCO, Thermo Scientific 89883) that had been equilibrated with the same assay buffer, and the protein concentration was determined with the Bradford assay.

The dephosphorylated KaiC (3.5 μM) was incubated at 30°C with 1.5 μM of either KaiA^WT^, KaiA^D52H^, or KaiA^V131A^ in the assay buffer containing either (i) 10 mM ATP only (= “100% ATP”), or (ii) 5 mM ATP and 5 mM ADP (= “50% ATP”). A portion of the phosphorylation reaction was withdrawn, mixed with an equal volume of 2x SDS-PAGE sample buffer, and stored at −20°C. Each sample was analyzed by SDS-PAGE to quantify the phosphorylation states of KaiC.^32,57^ To assess the effect of oxidation on KaiC autokinase activity, the reactions were treated with various concentrations of 2,3-dimethoxy-5-methyl-*p*-benzoquinone (Q_0_, Sigma-Aldrich D9150).

### *In vitro* KaiABC reaction monitored by fluorescence anisotropy

The *in vitro* oscillations were monitored as described by Heisler *et al.*^58^ with minor modifications. The *in vitro* reactions contained 3.5 μM KaiC, 1.5 μM KaiA^WT^, KaiA^D52H^, or KaiA^V131A^, 3.4 μM KaiB, and 0.1 μM of 6-iodoacetamidofluroescein (6IAF)-labeled KaiB^K25C^ in 20 mM Tris-HCl, pH 8.0, 150 mM NaCl, 5 mM MgCl_2_ and 10 mM ATP. The reactions were treated with 0.625, 1.25, 2.5, or 5 μM Q_0_. The reactions (30 or 100 μL per well) were added into the wells of a 384-well microplate (Greiner 781900). The microplate was sealed with a polyolefin film (Thermo Scientific 235307), and incubated at 30°C in a multimode microplate reader (Tecan Spark). Fluorescence anisotropy was measured every 15 min on the plate reader using polarized 485 nm (20 nm bandwidth) as excitation, with the emission measured at 535 nm (20 nm bandwidth filter) using a 510 nm dichroic splitter. Periods and amplitudes of the *in vitro* oscillations were determined by the Wavelet-based time-frequency analysis using pyBOAT.^59^

### Calculation of Q_10_

Q_10_ is a coefficient to express how much the rate of a reaction or biological process changes as the temperature changes. It measures how much the rate changes for every 10-degree Celsius increase in temperature. A Q_10_ value of exactly 1.0 means that the reaction is temperature independent, while a Q_10_ value greater than 1.0 means that the reaction accelerates as the temperature is increased (and conversely, a Q_10_ value less than 1.0 means that the rate *slows down* as the temperature is increased). For intervals of temperature changes that are exactly 10 Celsius degrees, Q_10_ is calculated as:

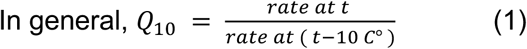

When calculating Q_10_ from data in which the temperature differential was not exactly 10 C°, the temperature coefficient (*Q*_10_) was calculated by:

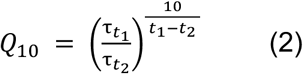

where 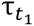 is period at temperature *t*_1_, and 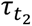 is period at temperature *t*_2_.

For period data at multiple different temperatures, temperature coefficient (*Q*_10_) was estimated by fitting equation (equation 3) to a set of experimental data points:

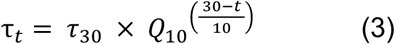

where *τ_t_* is period at temperature *t*. The fitting extracts temperature coefficient (*Q*_10_) and estimated period at 30°C (*τ*_30_).

### Estimation of cellular redox by measurement of chlorophyll fluorescence (in cyanobacteria)

In cyanobacteria, photosynthesis and respiration share a common plastoquinone (PQ) pool.^28^ As shown in Supplementary Fig. 1, the pool is reduced by Photosystem II (PSII), Photosynthetic Complex I, Succinate Dehydrogenase (SDH), and Proton Gradient Regulator 5 (PGR5). The pool is oxidized by cytochrome (cyt) b_6_f and various alternative oxidases such as cyt bd and PQ terminal oxidase (PTOX). The electron source or sink for each of these complexes is soluble, thus connecting the cytosolic and membrane redox states. PSII contains chlorophyll molecules that are excited by visible light and may relax via fluorescence. When the PQ pool is in a reduced state, the yield of PSII chlorophyll fluorescence is high. When the PQ pool is in an oxidized state, the yield of PSII chlorophyll fluorescence is low. This variable chlorophyll fluorescence signal provides a nonintrusive method to monitor intracellular redox in cyanobacteria.

To measure chlorophyll fluorescence, cyanobacterial strains were maintained on BG-11 plates with 1.5% (w/v) agar and without sodium thiosulfate at 30°C with 1% CO_2_ in air at 50 µmol photons m^−2^ s^−1^ cool white light. To prepare spot assays, cells on plates were subjected to darkness for 12 hours at 30°C and ambient CO_2_. Plates were maintained at 30°C and ambient CO_2_ thereafter. Following the first dark treatment, the plates were then subjected to 12 hours of light using either 50 µmol photons m^−2^ s^−1^ of cool white LED lighting, or 35 µmol photons m^−2^ s^−1^ of red (*λ*_max_ = 630 nm) or blue (*λ*_max_ = 462 nm) LED lighting. About halfway through this period, an aliquot of cells was removed from the plate, resuspended in liquid BG-11 media, and chlorophyll was quantified spectrophotometrically from a methanol extract. The suspension was normalized to 1.25 µg Chl µL^-1^ and 10 µL (12.5 µg chlorophyll) was used to spot fresh BG-11 plates containing no additives (control), DMSO (vehicle control), 50 µM VU661, 50 µM VU164, or 30 µM phenazine. These plates completed the 12-hour light period and were then subjected to a second 12-hour dark treatment. Plates under LL, RR, or BB conditions were measured after 24 hours using a fluorescence imager (FluorCam 800MF, Photon Systems Instruments, Brno, Czech Republic) fitted with a thermostat-controlled base. Actinic light at 35 µmol photons m^−2^ s^−1^ was provided by LED panels inside the imager (cool white, red (*λ*_max_ = 627 nm), or blue (*λ*_max_ = 447 nm). Chlorophyll fluorescence was quantified using non-actinic measuring flashes in the presence (F’) or absence (F_o_’) of actinic light and following a saturating flash (F_m_’). The fraction of open Photosystem II centers and thus the relative redox state of the PQ pool was estimated using the parameter q_L_^60^ defined as:

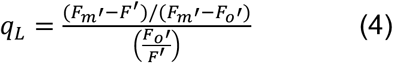

### Estimation of cellular redox by optical redox ratio (in mammalian cells)

Mammalian U2OS cells were seeded into 4-sector-glass bottom dishes (D35C4-20-1.5-N, Cellvis) and cultured with DMEM + 10 % fetal bovine serum, and 100 I.U. mL^-1^ Penicillin, 100 μg mL^-1^ Streptomycin for 24 h prior to the imaging experiment. On the day of the experiment, the culture medium was removed and replaced with this imaging buffer solution: Hank’s Balanced Salt Solution (Gibco 14025134), 25 mM glucose, 0.1% (w/v) bovine serum albumin; the solution is referred to as “vehicle” in the figures and in the legends. We used two-photon excitation spectral imaging to simultaneously excite the fluorescence of NADH and FAD from live U2OS cells, and we resolved the two emissions by linear unmixing. Images were acquired using a microscope LSM880 (Zeiss) with a 63X oil objective (Plan-Apochromat 63x/1.40 Oil DIC M27, Zeiss) coupled to a tunable infrared laser for multi-photon excitation (Chamaeleon Discovery, Coherent), and a stage-top incubator to maintain 37 °C, 5% CO_2_, 100% humidity (Pecon). We used an external temperature probe, placed in a sector adjacent to the one containing the cells being imaged, to confirm that the cells were exposed to the desired temperature. The imaging settings were the following: excitation wavelength 710 nm, emission bandpass for the spectral detector 410 nm-624 nm, image size 1024 × 1024 pixels, pixel size 0.22 μm, pixel dwell time 16.2 μs. The laser power output was 1910 mW and it was attenuated to 1.8%; these settings provided adequate fluorescence intensity without saturating the detector and without photodamage to the cells. Each spectral stack was processed using the ZEN software (Zeiss) to extract individual fluorescent components by linear unmixing. NADH salt, FAD salt (Sigma), and all the tested compounds were dissolved in vehicle to acquire their reference fluorescence spectra for linear unmixing. Compound VU661 has a fluorescence emission centered around 440 nm that could be distinguished from the partially overlapping emission of NADH centered at 460 nm; compound VU164 has a very dim fluorescence that was negligible when compared to the endogenous cellular emissions. The segmentation and the analysis of the unmixed images was done using Fiji as follows: the NADH image was used to define the region of interest (ROI) attributed to mitochondria, and to manually segment the nuclei. First, we removed any saturated pixel from the image using the Yen automatic threshold, then we applied a Fast Fourier Transform Band Pass Filter with a window between 3 and 35 pixels; last, we applied the Triangle auto threshold function to segment the ROI containing the mitochondria. The ROI containing the nuclei were drawn manually on each cell, as the nucleus was clearly visible as a dark circular region. We used the sum of the NADH and the FAD images to define the ROI of the entire cells by applying the Median Filter with radius 6 and the applying the Percentile auto threshold. The area excluded from the mitochondria and from the nuclei ROIs was assigned to the cytoplasm ROI. After the 3 ROIs were created, we built an optical redox ratio (ORR) image where each pixel value is determined by the following formula ORR = FAD/(NADH+FAD). The intensity measurements were performed on the unfiltered NADH and FAD images, and on the ORR images, to determine the mean pixel intensity in each ROI.

### Assay of energy charge in cyanobacterial cells

Energy charge (E.C.) was calculated from ATP and ADP levels as (ATP)/(ATP + ADP) using a methodology adapted from Pattanayak *et al.*^23^ Cyanobacterial cultures were grown on BG-11 petri dishes and harvested by pipetting 5 mL of liquid BG-11 medium ^47^ onto the agar plates. Using a plastic L-shaped spreader, cells were gently resuspended into the liquid medium, and 3 mL of the cell/medium suspension was quickly added to a 15 mL test tube containing 750 μL ice-cold 1M hydrochloric acid, vortexed, and placed on ice. One mL of the cell/medium/acid suspension was removed and used for measurement of OD_750_ (which varied from preparation to preparation, but was generally ∼0.6). To permeabilize the cells by freeze/thaw, the remaining 2 mL in the tubes was placed at −80°C for 1 h and thereafter thawed by incubation in a 65°C water bath for 10 min. The pH of the suspension was neutralized with 600 μL of 1 M KOH, 0.5 M Tris, 0.5 M KCl, plus 1.4 mL of water (total volume 2 mL) and centrifuged at 4000 rpm for 10 min. The supernatant was collected from each tube and stored at −80°C.

On the day of assay, the supernatant samples were thawed on ice. For each timepoint/treatment, a “ATP-only” sample and a “[ATP+ADP]” sample was generated. For measuring the “ATP-only” sample, 220 μL of the supernate sample was added to 352 μL of 25 mM KCl, 50 mM MgSO_4_ and 100 mM HEPES pH 7.4 (LK buffer) in a 0.6 mL microcentrifuge tube. To generate the “[ATP+ADP]” sample, ADP was converted to ATP for assay by combining 220 μL of the supernatant sample with 330 μL LK buffer, 11 μL of 100 mM phosphoenolpyruvate (PEP), and 11 μL of 250 U mL^-1^ type II pyruvate kinase from rabbit muscle (Sigma-Aldrich). The “[ATP+ADP]” samples were incubated at 37°C in a water bath for 1 h to convert ADP to ATP and then the pyruvate kinase reaction was stopped at 96°C (10 min). The “ATP-only” and the “[ATP+ADP]” samples were brought to room temperature and assayed for ATP levels with the firefly luciferase assay.^23^ Standard curves for ATP levels were based on standards consisting of 300, 150, 75, 37.5, 18.75, 9.375, and 0 nM ATP that were made by serial dilution in LK buffer, and 20 μL of each standard was added to a well in a 96-well plate that contained 210 μL of LK buffer. The “ATP-only” and the “[ATP+ADP]” wells contained 20 μL of the respective sample and as an internal standard, each sample well contained in addition 20 μL of an ATP standard (either 300, 150, 75, or 0 nM ATP) plus 190 μL of LK buffer. A working solution of luciferin (Sigma-Aldrich) and firefly luciferase (Sigma-Aldrich) in LK buffer was made fresh for each assay and kept on ice until the assay was conducted (working solution: 60 μg mL^-1^ firefly luciferase and 1.67 mM firefly luciferin). In a PolarStar Optima plate reader, 30 μL of the luciferin/luciferase solution was rapidly injected into each well, and the luminescence signal emanating from the well was immediately recorded for 10 sec. Concentrations of ATP were determined by comparison with the linear regression from the standard curve, and the internal standards within each of the samples confirmed the validity of each assay. The ATP values for the “ATP-only” and the “[ATP+ADP]” samples were then used in the equation (ATP)/(ATP + ADP) to calculate the energy charge.

### Models and simulations

#### Mathematical model for the KaiC post-translational oscillator coupled to a transcriptional-translational feedback loop (PTO/TTFL)

We developed a mathematical model of the phosphorylation cycle of KaiC coupled to a transcriptional-translational feedback loop (TTFL; Fig. 3c). We adopt the mathematical model proposed by Rust *et al.*^61^ and Teng *et al.*^62^ for the description of the phosphorylation cycle of KaiC. The model includes four different KaiC phosphorylation states, *U*, *T*, *D*, & *S* to label the levels of: (i) unphosphorylated KaiC (U), (ii) KaiC phosphorylated at T432 (T), (iii) KaiC phosphorylated at both T432 and S431 (D, or also referred to as “ST”), and (iv) KaiC phosphorylated at S431 (S) (Fig. 3c). In cyanobacteria, phosphorylated KaiC regulates its own transcription through the activation of the global transcriptional factor RpaA *in vivo* (Fig. 3c). We model this by modifying the transcriptional regulation term in Teng *et al.*^62^ In our model, the doubly phosphorylated KaiC (*D*) activates the linker kinase, SasA, which phosphorylates RpaA. The phosphorylated RpaA activates the transcription of the *kaiC* gene. Newly translated KaiC protein enters the phosphorylation cycle in its unphosphorylated (U) state. We assume a linear degradation of KaiC protein with the same degradation rate *V_d_* among the different phosphorylation states. In the mathematical model, the total KaiA levels are constant over time. The *kaiC* mRNA levels both (i) indicate the action of the TTFL back onto the IVO/PTO, and also (ii) serve as a proxy for global circadian transcription.^20,33^ The time evolution of the levels of each KaiC phosphorylation state and *kaiC* mRNA levels (*M*) is described as:

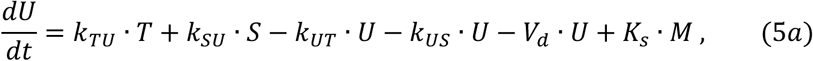

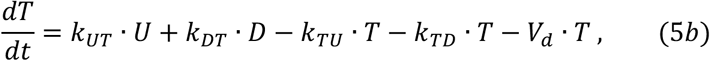

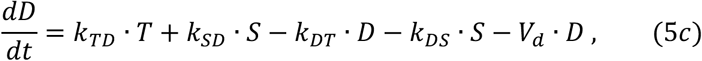

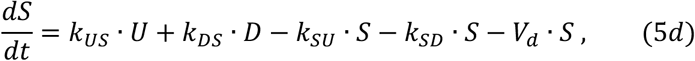

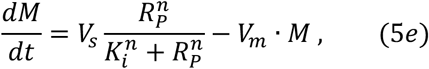

where *k_XY_* (*X*, *Y* ∈ {*U*, *T*, *D*, *S*}) is a KaiA-dependent reaction rate, *K_s_* is the translation rate of KaiC protein and *V_s_* is the maximum transcription rate of *kaiC* mRNA. The translation of KaiC in Equation (5a) is proportional to *kaiC* mRNA levels *M*. In the first term of Equation (5e), we use a Hill function for the transcriptional regulation of *kaiC* by phosphorylated RpaA (*R_p_*). The *n* in Equation (5e) is the Hill coefficient and *K_i_* is a threshold constant determining the RpaA levels for the half maximum value of transcription. The *kaiC* mRNA is degraded at the rate *V_m_*.

The reaction rates *k_XY_* in Equations (5a-d) depend on the free KaiA levels *A*(*S*) as in Rust *et al.*^61^

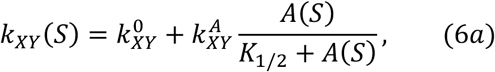

where:

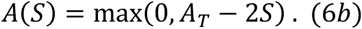

The first and second terms of Equation (6a) are the KaiA-independent and KaiA-dependent reaction rates, respectively. 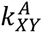 is the maximum reaction rate at saturating levels of free KaiA and *K*_1/2_. is the dissociation constant of free KaiA from KaiC. Equation (6b) represents the sequestration of KaiA by KaiC in its *S* form. *A_T_* in Equation (6b) is the total KaiA levels.

We describe the time evolution of phosphorylated RpaA levels *R_p_* (RpaA-P in (Fig. 3c) by the following differential equation:

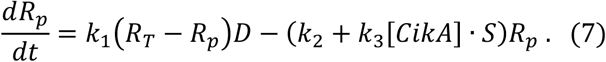

The first term of Equation (7) is the phosphorylation of RpaA by doubly phosphorylated KaiC (*D*). We assume that the total RpaA levels (*R_T_*) are constant over time. Therefore, *R_T_* – *R_p_* represents unphosphorylated RpaA levels. The second term of Equation (7) is the dephosphorylation of RpaA: *k*_2_ is the basal dephosphorylation rate of RpaA-P and [*CikA*] is the CikA levels. Based on a previous experimental study ^63^, we simulate that CikA promotes the dephosphorylation of RpaA-P by interacting with KaiC in its *S* form.

#### Mathematical model for the *in vitro* KaiC post-translational oscillator (IVO/PTO)

The *in vitro* phosphorylation cycle of KaiC was simulated by setting *K_s_* = 0 and *V_d_* = 0 in the *in vivo* model {i.e., Equation (5)}. We also changed the *K*_1/2_ values to consider the difference in KaiA activity or KaiA levels between *in vivo* and *in vitro* conditions (Extended Data Table 1). Because the *in vitro* total KaiC levels [KaiC] = *U* + *T* + *D* + *S* remain constant, we solved the differential equations for *T*, *D*, and *S* in Equation (5), and calculated *U* by the relation *U* = [KaiC] − (*T* + *D* + *S*) in numerical simulations.

#### Description of KaiA mutants and modeling the “oxidization” condition

To model the influence of mutated KaiA, we increased the value of *K*_1/2._ in Equation (6a) from the wild-type value (Extended Data Table 2). Our experimental data indicates that both KaiA activity and CikA levels are reduced under oxidizing redox conditions. In the case of KaiA activity, the oxidizing condition of Q_0_ results in a greater reduction of the KaiA-stimulation of KaiC autophosphorylation in KaiA^D52H^ and KaiA^V131A^ than in KaiC^WT^ (Fig. 3e & Extended Data Fig. 5d,e), as well as a greater impact upon amplitude and damping of the PTO with the KaiA mutants (Extended Data Fig. 5a-c). In the case of CikA, the Golden lab reported that the oxidizing reagent DBMIB reduced CikA levels.^36^ We therefore tested an oxidizing condition (30 μM phenazine), and confirmed that CikA levels significantly decreased after the addition of phenazine (Extended Data Fig. 6). Therefore, we modeled oxidizing redox conditions by increasing *K*_1/2_ in Equation (6a) and decreasing CikA levels in Equation (7) from the standard (non-oxidizing) condition (Extended Data Table 2). Because the reduction of KaiA activity by oxidization is greater in the KaiA mutants than wildtype (Fig. 3e & Extended Data Fig. 5d,e), we assign a larger change in *K*_1/2_ in KaiA mutants than in KaiA^WT^ in these simulations.

#### Numerical methods

We numerically solved (Equations 5)-(7) by using the fourth-order Runge-Kutta method implemented with MATLAB 2024b. Initial conditions of simulations are (*U, T, D, S, M, R_p_*) = (2.23, 0.83, 0.31, 0.48, 0.037, 0) for the *in vivo* model and (*T, D, S*) = (0, 1, 0.2) for the *in vitro* model. The values of parameters for simulations are listed in Extended Data Tables 1 and 2. The values of parameters for phosphorylation cycle of KaiC are based on those in Rust *et al.*^61^ Values of other parameters involved in the *kaiC* TTFL were chosen so that the period of the transcription rhythm of *kaiC* in wild type was in the circadian range.

## Acknowledgments

We thank multiple colleagues for discussions, assistance, and wisdom regarding this project, especially Achim Kramer, Hans Peter Herzel, Marta del Olmo Somolinos, Jamey Young, Yong-Ick Kim, Ian Dew, Sara Chen, Emily Bishop, Caroline McCool, and Kevin Kelly. This work was supported by USA National Institutes of Health grant R37 GM067152 (CHJ), USA National Institutes of Health grant R01 DK123301 (DWP), and USA Department of Energy grant DE-SC0025359 (DV).

## Author contributions

Conceptualization: YX, TM, CHJ; Methodology: YX, TM, HM, AU, KT, YK, KU, DWP, DV, CHJ; Investigation: YX, TM, HM, AU; Visualization: YX, TM, HM, AU, YK, KU, DV, CHJ; Mathematical modeling: YK, KU; Funding acquisition: CHJ, DWP, DV; Project administration: CHJ; Supervision: CHJ, DV, DWP, KU, SN; Writing – original draft: YX, TM, YK, KU, DV, CHJ; Writing – review & editing: YX, TM, HM, AU, KT, YK, KU, DWP, DV, CHJ.

## Data and materials availability

The authors declare that all data supporting the results of this study are available within the main text or the supplementary materials.

## References

1 Johnson, C. H., Elliott, J. A., Foster, R. G., Honma, K.-I. & Kronauer, R. Chapter 3: Fundamental properties of circadian rhythms. in Chronobiology: Biological Timekeeping (eds Dunlap, J. C., Loros, J. J. & DeCoursey, P. J.) 66–105 (Sinauer Associates, Sunderland, Mass, 2004).

2. Ouyang, Y., Andersson, C. R., Kondo, T., Golden, S. S. & Johnson, C. H. Resonating circadian clocks enhance fitness in cyanobacteria. Proc Natl Acad Sci U S A 95, 8660–8664 (1998).

3. Jabbur, M. L., Dani, C., Spoelstra, K., Dodd, A. N. & Johnson, C. H. Evaluating the adaptive fitness of circadian clocks and their evolution. J Biol Rhythms 39, 115–134 (2024).

4. Pittendrigh, C. S. & Caldarola, P. C. General homeostasis of the frequency of circadian oscillations. Proc Natl Acad Sci U S A 70, 2697–2701 (1973).

5. Brown, S. A., Zumbrunn, G., Fleury-Olela, F., Preitner, N. & Schibler, U. Rhythms of mammalian body temperature can sustain peripheral circadian clocks. Curr Biol 12, 1574–1583 (2002).

6. Hastings, J. W. & Sweeney, B. M. On the mechanism of temperature independence in a biological clock. Proc Natl Acad Sci U S A 43, 804–811 (1957).

7. Mehra, A. et al. A role for casein kinase 2 in the mechanism underlying circadian temperature compensation. Cell 137, 749–760 (2009).

8. Terauchi, K. et al. ATPase activity of KaiC determines the basic timing for circadian clock of cyanobacteria. Proc Natl Acad Sci U S A 104, 16377–16381 (2007).

9. Zhou, M., Kim, J. K., Eng, G. W. L., Forger, D. B. & Virshup, D. M. A Period2 phosphoswitch regulates and temperature compensates circadian period. Mol Cell 60, 77–88 (2015).

10. Stevenson, E.-L., Mehalow, A. K., Loros, J. J., Kelliher, C. M. & Dunlap, J. C. A compensated clock: Temperature and nutritional compensation mechanisms across circadian systems. Bioessays 47, e202400211 (2025).

11. Rutter, J., Reick, M. & McKnight, S. L. Metabolism and the control of circadian rhythms. Annu Rev Biochem 71, 307–331 (2002).

12. Kohsaka, A. et al. High-fat diet disrupts behavioral and molecular circadian rhythms in mice. Cell Metab 6, 414–421 (2007).

13. Bass, J. Circadian topology of metabolism. Nature 491, 348–356 (2012).

14. Laothamatas, I., Rasmussen, E. S., Green, C. B. & Takahashi, J. S. Metabolic and chemical architecture of the mammalian circadian clock. Cell Chem Biol 30, 1033–1052 (2023).

15. Asher, G. & Schibler, U. Crosstalk between components of circadian and metabolic cycles in mammals. Cell Metab 13, 125–137 (2011).

16. Diamond, S., Jun, D., Rubin, B. E. & Golden, S. S. The circadian oscillator in *Synechococcus elongatus* controls metabolite partitioning during diurnal growth. Proc Natl Acad Sci U S A 112, E1916–1925 (2015).

17. Johnson, C. H. & Egli, M. Metabolic compensation and circadian resilience in prokaryotic cyanobacteria. Annu Rev Biochem 83, 221–247 (2014).

18. Chance, B. & Thorell, B. Fluorescence measurements of mitochondrial pyridine nucleotide in aerobiosis and anaerobiosis. Nature 184, 931–934 (1959).

19. Pittendrigh, C. S., Caldarola, P. C. & Cosbey, E. S. A differential effect of heavy water on temperature-dependent and temperature-compensated aspects of circadian system of *Drosophila pseudoobscura*. Proc Natl Acad Sci U S A 70, 2037–2041 (1973).

20. Circadian Rhythms in Bacteria and Microbiomes. (eds Johnson, C. H. & Rust, M. J., Springer International Publishing, Cham, 2021). doi:10.1007/978-3-030-72158-9.

21. Ishiura, M. et al. Expression of a gene cluster *kaiABC* as a circadian feedback process in cyanobacteria. Science 281, 1519–1523 (1998).

22. Kondo, T. et al. Circadian clock mutants of cyanobacteria. Science 266, 1233–1236 (1994).

23. Pattanayak, G. K., Lambert, G., Bernat, K. & Rust, M. J. Controlling the cyanobacterial clock by synthetically rewiring metabolism. Cell Rep 13, 2362–2367 (2015).

24. Rocheleau, J. V., Head, W. S. & Piston, D. W. Quantitative NAD(P)H/flavoprotein autofluorescence imaging reveals metabolic mechanisms of pancreatic islet pyruvate response. J Biol Chem 279, 31780–31787 (2004).

25. Wood, T. L. et al. The KaiA protein of the cyanobacterial circadian oscillator is modulated by a redox-active cofactor. Proc Natl Acad Sci U S A 107, 5804–5809 (2010).

26. Fang, M., Chavan, A. G., LiWang, A. & Golden, S. S. Synchronization of the circadian clock to the environment tracked in real time. Proc Natl Acad Sci U S A 120, e2221453120 (2023).

27. Kelly, K. P. et al. Screen for small-molecule modulators of circadian rhythms reveals phenazine as a redox-state modifying clockwork tuner. ACS Chem Biol 17, 1658–1664 (2022).

28. Mullineaux, C. W. Co-existence of photosynthetic and respiratory activities in cyanobacterial thylakoid membranes. Biochim Biophys Acta 1837, 503–511 (2014).

29. Calzadilla, P. I. & Kirilovsky, D. Revisiting cyanobacterial state transitions. Photochem Photobiol Sci 19, 585–603 (2020).

30. Tanaka, K. et al. The endogenous redox rhythm is controlled by a central circadian oscillator in cyanobacterium *Synechococcus elongatus* PCC7942. Photosynth Res 142, 203–210 (2019).

31. Lu, Y. et al. Regulation of the cyanobacterial circadian clock by electrochemically controlled extracellular electron transfer. Angew Chem Int Ed Engl 53, 2208–2211 (2014).

32. Nakajima, M. et al. Reconstitution of circadian oscillation of cyanobacterial KaiC phosphorylation *in vitro*. Science 308, 414–415 (2005).

33. Chavan, A. G. et al. Reconstitution of an intact clock reveals mechanisms of circadian timekeeping. Science 374, (2021).

34. Kim, Y.-I., Dong, G., Carruthers, C. W., Golden, S. S. & LiWang, A. The day/night switch in KaiC, a central oscillator component of the circadian clock of cyanobacteria. Proc Natl Acad Sci U S A 105, 12825–12830 (2008).

35. Mori, T. et al. Revealing circadian mechanisms of integration and resilience by visualizing clock proteins working in real time. Nat Commun 9, 3245 (2018).

36. Ivleva, N. B., Gao, T., LiWang, A. C. & Golden, S. S. Quinone sensing by the circadian input kinase of the cyanobacterial circadian clock. Proc Natl Acad Sci U S A 103, 17468–17473 (2006).

37. Ruby, N. F., Burns, D. E. & Heller, H. C. Circadian rhythms in the suprachiasmatic nucleus are temperature-compensated and phase-shifted by heat pulses *in vitro*. J Neurosci 19, 8630–8636 (1999).

38. Izumo, M., Johnson, C. H. & Yamazaki, S. Circadian gene expression in mammalian fibroblasts revealed by real-time luminescence reporting: temperature compensation and damping. Proc Natl Acad Sci U S A 100, 16089–16094 (2003).

39. Skala, M. C. et al. In vivo multiphoton microscopy of NADH and FAD redox states, fluorescence lifetimes, and cellular morphology in precancerous epithelia. Proc Natl Acad Sci U S A 104, 19494–19499 (2007).

40. Goto, K., Laval-Martin, D. L. & Edmunds, L. N. Biochemical modeling of an autonomously oscillatory circadian clock in *Euglena*. Science 228, 1284–1288 (1985).

41. Wang, T. A. et al. Circadian rhythm of redox state regulates excitability in suprachiasmatic nucleus neurons. Science 337, 839–842 (2012).

42. del Olmo Somolinos, M. *Searching for Order in Body Clocks. Circadian Rhythms and Redox Balance*. (Logos Verlag Berlin, Berlin, 2021). doi:10.30819/5406.

43. Shinohara, Y. et al. Temperature-sensitive substrate and product binding underlie temperature-compensated phosphorylation in the clock. Mol Cell 67, 783–798.e20 (2017).

44. Isojima, Y. et al. CKIε/δ-dependent phosphorylation is a temperature-insensitive, period-determining process in the mammalian circadian clock. Proc Natl Acad Sci U S A 106, 15744–15749 (2009).

45. O’Neill, J. S. & Reddy, A. B. Circadian clocks in human red blood cells. Nature 469, 498–503 (2011).

46. O’Neill, J. S. et al. Circadian rhythms persist without transcription in a eukaryote. Nature 469, 554–558 (2011).

47. Bustos, S. A. & Golden, S. S. Expression of the *psbDII* gene in *Synechococcus sp.* strain PCC 7942 requires sequences downstream of the transcription start site. J Bacteriol 173, 7525–7533 (1991).

48. Cirino, P. C., Mayer, K. M. & Umeno, D. Generating mutant libraries using error-prone PCR. Methods Mol Biol 231, 3–9 (2003).

49. Johnson, C. H. & Xu, Y. The Decade of Discovery: How *Synechococcus elongatus* Became a Model Circadian System 1990–2000, Chapter 4. in Bacterial Circadian Programs (eds Ditty, J. L., Mackey, S. R. & Johnson, C. H.) 63–86 (Springer, Berlin, Heidelberg, 2009). doi:10.1007/978-3-540-88431-6_4.

50. Xu, Y. et al. Non-optimal codon usage is a mechanism to achieve circadian clock conditionality. Nature 495, 116–120 (2013).

51. Xu, Y. et al. Intramolecular regulation of phosphorylation status of the circadian clock protein KaiC. PLoS One 4, e7509 (2009).

52. Nishio, K. et al. Extracellular electron transfer across bacterial cell membranes via a cytocompatible redox-active polymer. Chemphyschem 14, 2159–2163 (2013).

53. Zhang, E. E. et al. A genome-wide RNAi screen for modifiers of the circadian clock in human cells. Cell 139, 199–210 (2009).

54. Izumo, M., Sato, T. R., Straume, M. & Johnson, C. H. Quantitative analyses of circadian gene expression in mammalian cell cultures. PLoS Comput Biol 2, e136 (2006).

55. Yamazaki, S. & Takahashi, J. S. Real-time luminescence reporting of circadian gene expression in mammals. Methods Enzymol 393, 288–301 (2005).

56. Taylor, W., Wilson, S., Presswood, R. & Hastings, J. W. Circadian rhythm data collection with the Apple II microcomputer. J Interdiscip Cycle Res 13, 71–79 (1982).

57. Mori, T. et al. Elucidating the ticking of an *in vitro* circadian clockwork. PLoS Biol 5, e93 (2007).

58. Heisler, J., Chavan, A., Chang, Y.-G. & LiWang, A. Real-time *in vitro* fluorescence anisotropy of the cyanobacterial circadian clock. Methods Protoc 2, 42 (2019).

59. Schmal, C., Mönke, G. & Granada, A. E. Analysis of complex circadian time series data using wavelets. Methods Mol Biol 2482, 35–54 (2022).

60. Kramer, D. M., Johnson, G., Kiirats, O. & Edwards, G. E. New fuorescence parameters for the determination of Q_A_ redox state and excitation energy fluxes. Photosynth Res 79, 209 (2004).

61. Rust, M. J., Markson, J. S., Lane, W. S., Fisher, D. S. & O’Shea, E. K. Ordered phosphorylation governs oscillation of a three-protein circadian clock. Science 318, 809–812 (2007).

62. Teng, S.-W., Mukherji, S., Moffitt, J. R., de Buyl, S. & O’Shea, E. K. Robust circadian oscillations in growing cyanobacteria require transcriptional feedback. Science 340, 737–740 (2013).

63. Gutu, A. & O’Shea, E. K. Two antagonistic clock-regulated histidine kinases time the activation of circadian gene expression. Mol Cell 50, 288–294 (2013).

64. Qin, X., Byrne, M., Xu, Y., Mori, T. & Johnson, C. H. Coupling of a core post-translational pacemaker to a slave transcription/translation feedback loop in a circadian system. PLoS Biol 8, e1000394 (2010).

65. Abe, J. et al. Circadian rhythms. Atomic-scale origins of slowness in the cyanobacterial circadian clock. Science 349, 312–316 (2015).

