## Supplementary Information for "Circadian temperature compensation is intrinsically linked with metabolism and redox signaling"

### Extended Data Figs. 1 to 10

**Extended Data Fig. 1**

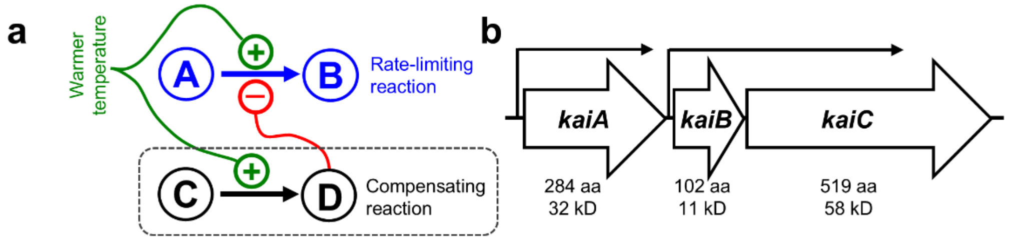


c

**High-throughput screening of a mutagenized *kai* library at 25°C from a single experiment**

| Mutant phenotype | Altered period (h) relative to wild type: < 20 | 20 ~ 25 | 27~ 35 | 35 ~ 45 | 45 ~ 55 | 55 ~ 65 | > 65 | Arrhy-thmicity | Low amplitude | Total # screened |
|---|---|---|---|---|---|---|---|---|---|---|
| # obtained | 1 | 78 | 63 | 17 | 8 | 5 | 2 | 262 | 14 | 9024 |

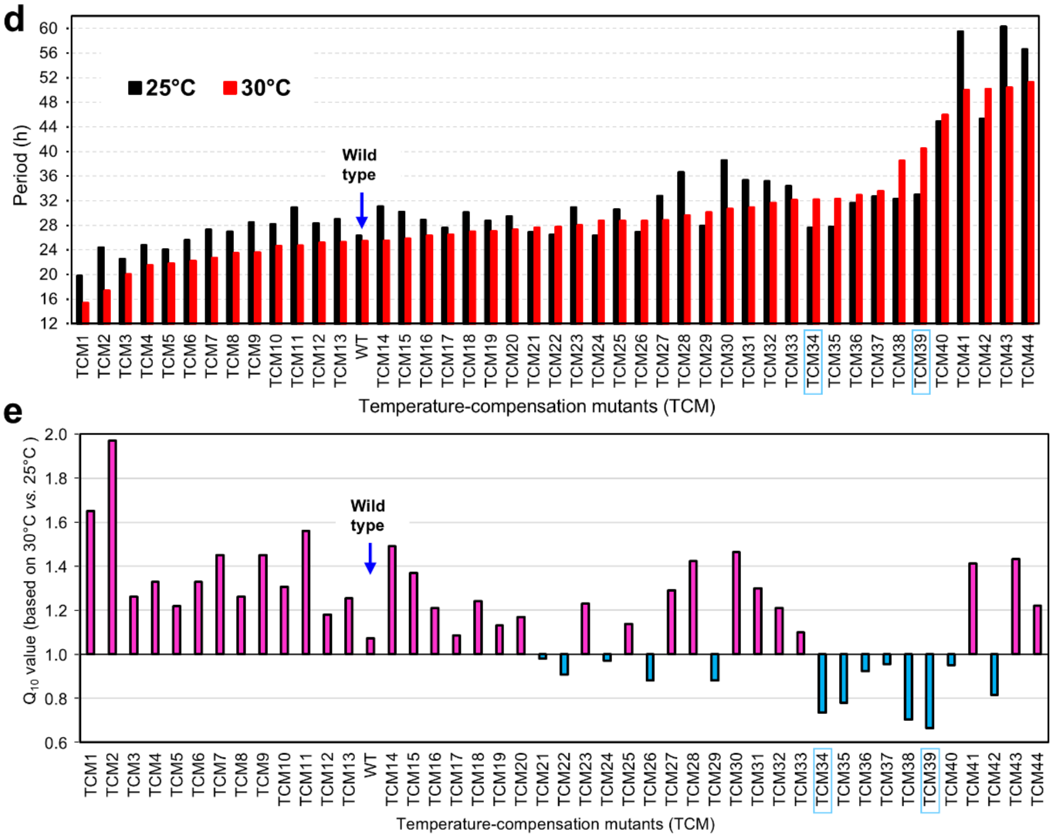


**Extended Data Fig. 1. Discovery of mutations in the *kaiABC* cluster that cause defects in circadian temperature compensation in cyanobacteria. a.** Hypothetical model of counterbalancing enzymatic reactions that was proposed to account for temperature compensation by Hastings & Sweeney in 1957.[6] Two reactions are involved, both of which separately have a $Q_{10} > 1.0$ (i.e., they speed up as the temperature increases). The reaction A → B is the rate-limiting reaction that determines circadian period, while C $\rightarrow$ D is the "compensating reaction." The product D inhibits the A $\rightarrow$ B reaction. If the two reactions are balanced appropriately, as the temperature increases, the C $\rightarrow$ D reaction slows down the temperature-induced acceleration of the A $\rightarrow$ B reaction, resulting in a final A $\rightarrow$ B reaction that is relatively unaffected by temperature changes. **b.** Diagram of the *kaiABC* cluster[21] used for error-prone PCR mutagenesis of *kaiABC* for screening for temperature compensation mutants at separate temperatures. **c.** A representative screening experiment at 25°C for temperature compensation mutants to illustrate the efficiency of the screening strategy. **d.** FRPs at 25°C versus 30°C genetic screening for temperature compensation mutants (TCMs). **e.** $Q_{10}$ values calculated from the FRPs at 25°C and 30°C for the TCM strains shown in panel **d**. $Q_{10}$ values below 1.0 are shown below the $Q_{10}$ = 1.0 line and plotted as inverted blue histograms. TCM34 and TCM 39 are the 2 key KaiA mutants V131A and D52H that are analyzed in this paper (here boxed in blue). Raw data of circadian luminescence rhythms for key strains analyzed in this paper are shown in Extended Data Fig. 10.

**Extended Data Fig. 2**

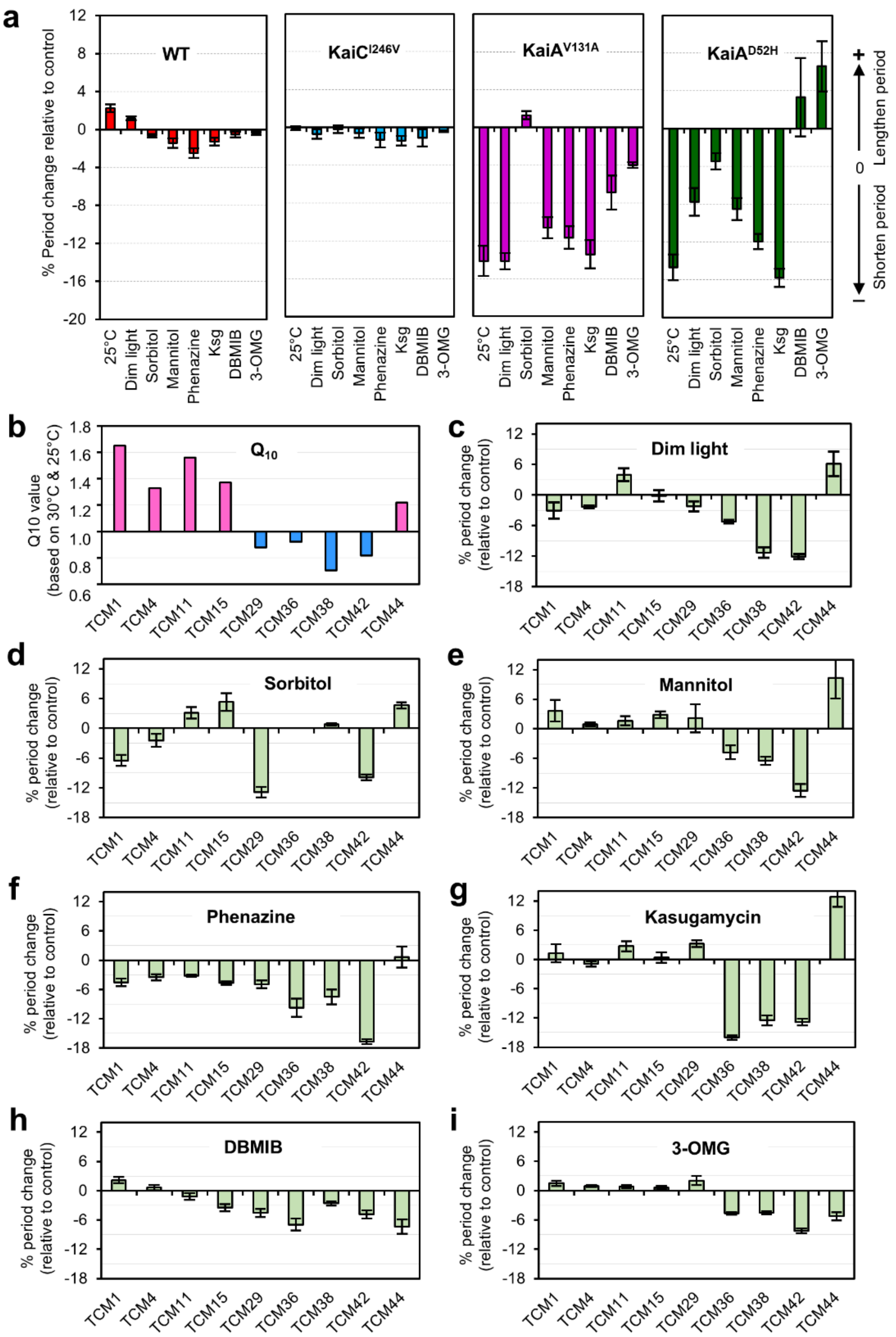


**Extended Data Fig. 2. "Metabolic conditions" affect the circadian FRP of TC-mutants but not of wild type cyanobacteria. a.** Metabolic perturbations significantly alter the FRP of *kaiA* TC mutants KaiA$^{V131A}$ and KaiA$^{D52H}$, but the FRP of the temperature-compensated WT and long-period mutant KaiC$^{I246V}$ strains are essentially unaffected. Sorbitol is a non-metabolizable sugar alcohol that functions here as a control for the metabolizable sugar alcohol mannitol. The data presented in this panel are the same as those appearing in Fig. 1d, except they are plotted here as "% period change," where negative values denote FRP shortening and positive values indicate FRP lengthening. **b.** $Q_{10}$ values of additional TCM mutant strains that harbor point mutations in either the *kaiA* or the *kaiC* genes (from Fig. 1d/e). **c** – **i.** FRP changes (plotted as "% period change" relative to corresponding controls) of the TCMs in panel b to conditions that perturb the metabolism of cyanobacterial cells: dim light of 10 µE $m^{-2}\cdot sec^{-1}$ (**c**), 3% sorbitol (**d**), 3% mannitol (**e**), 20 µM phenazine (**f**), 2 µg/ml kasugamycin (**g**), 30 µM DBMIB (**h**), and 0.2% 3-OMG (**i**), respectively. Error bars represent means ± S.D (n = 3).

**Extended Data Fig. 3**

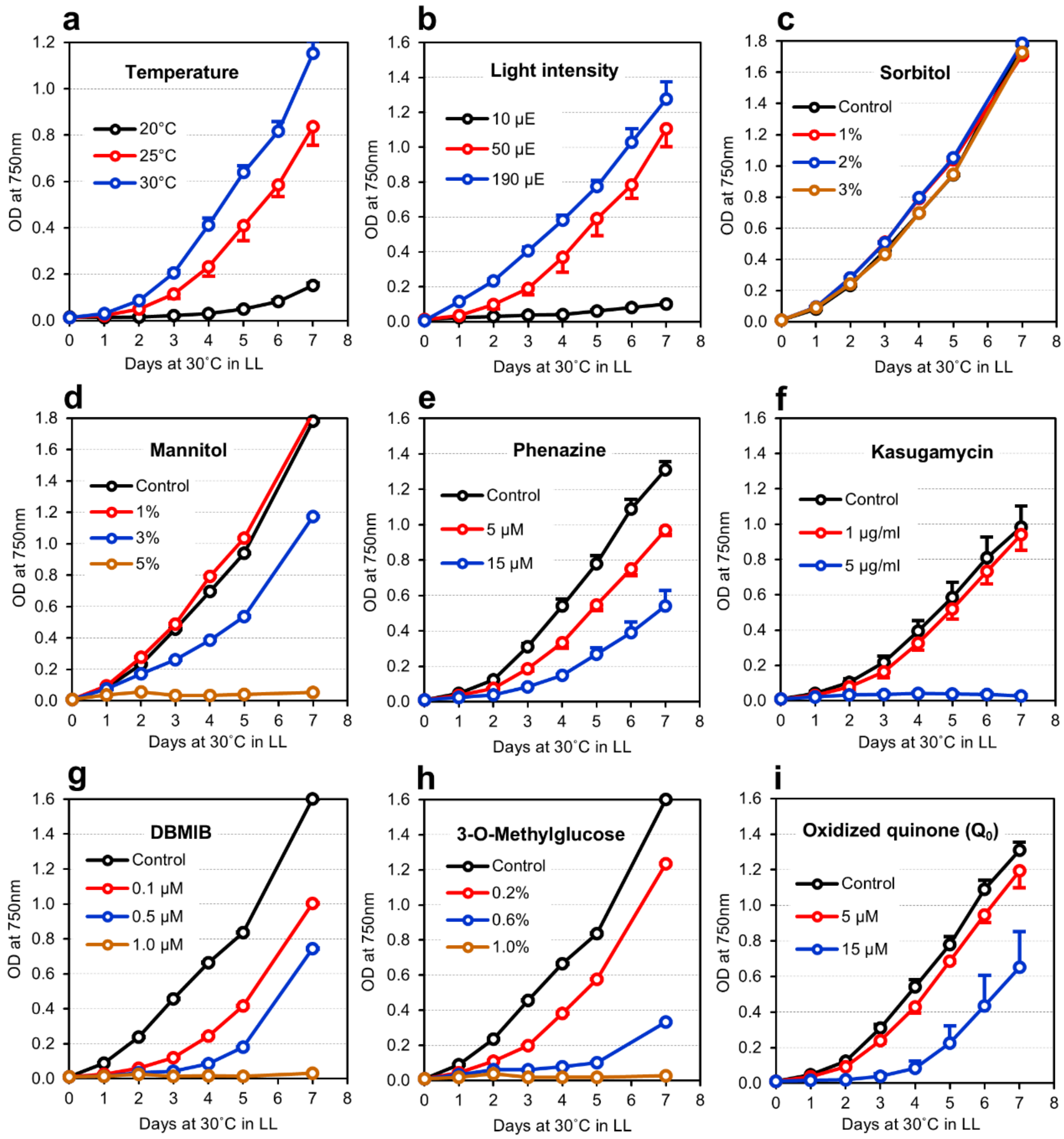


**Extended Data Fig. 3. Growth of wild-type cyanobacterial cultures under the metabolic conditions identified in Fig. 1d and Extended Data Fig. 2.** Growth of wild-type cyanobacterial cells in liquid cultures measured as the increase of light scattering at 750 nm ($O.D._{750nm}$) as a proxy for metabolic inhibition. The effects of different temperatures (**a**), light intensities as µE $m^{-2}\cdot sec^{-1}$ (**b**), and concentrations of treatments, some of which inhibit metabolism and growth (**c** – **i**) are as indicated.

**Extended Data Fig. 4**

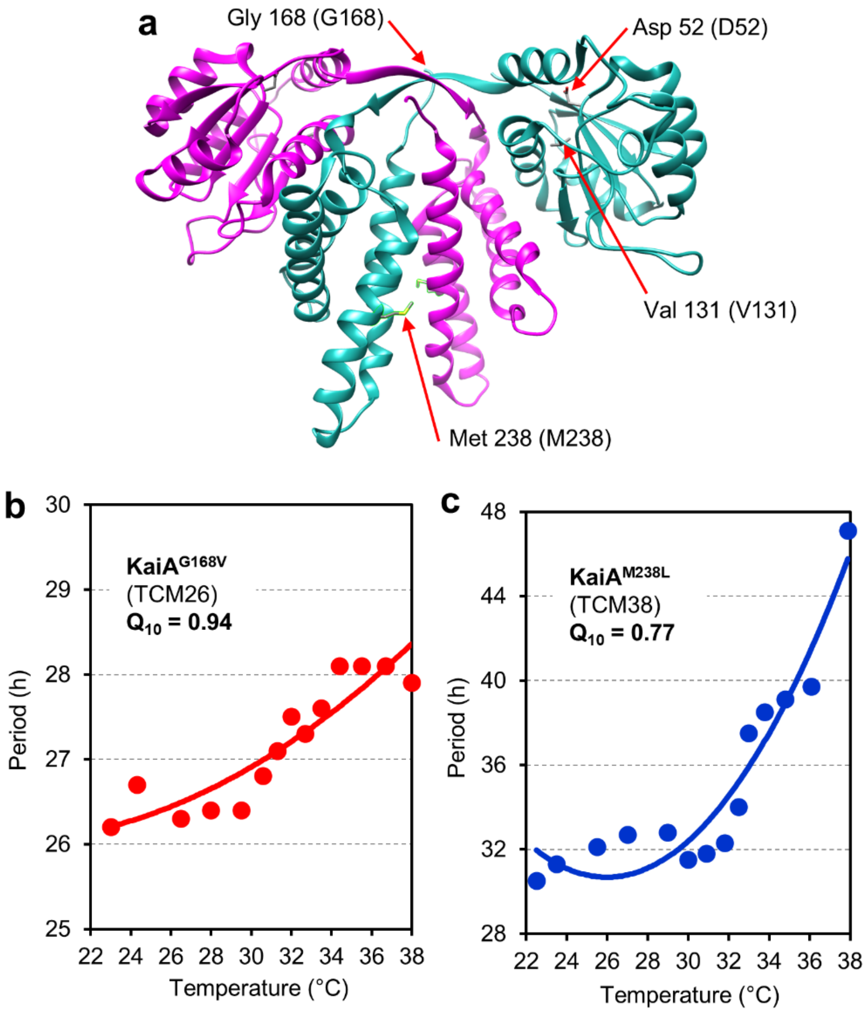


**Extended Data Fig. 4. Four point mutations in the *kaiA* gene alter temperature compensation properties.** $KaiA^{V131A}$ and $KaiA^{D52H}$ are shown in Fig. 1, but two additional point mutations in the *kaiA* gene are defective in temperature compensation with $Q_{10}$ values below 1.0. **a.** Locations of the G168V and M238L mutations along with the D52H and V131A mutations in the KaiA structure. **b & c.** Temperature dependency of FRPs and $Q_{10}$ values for the G168V (**b**) and M238L (**c**) mutants over the temperature range 22°C - 38°C.

#### Extended Data Fig. 5

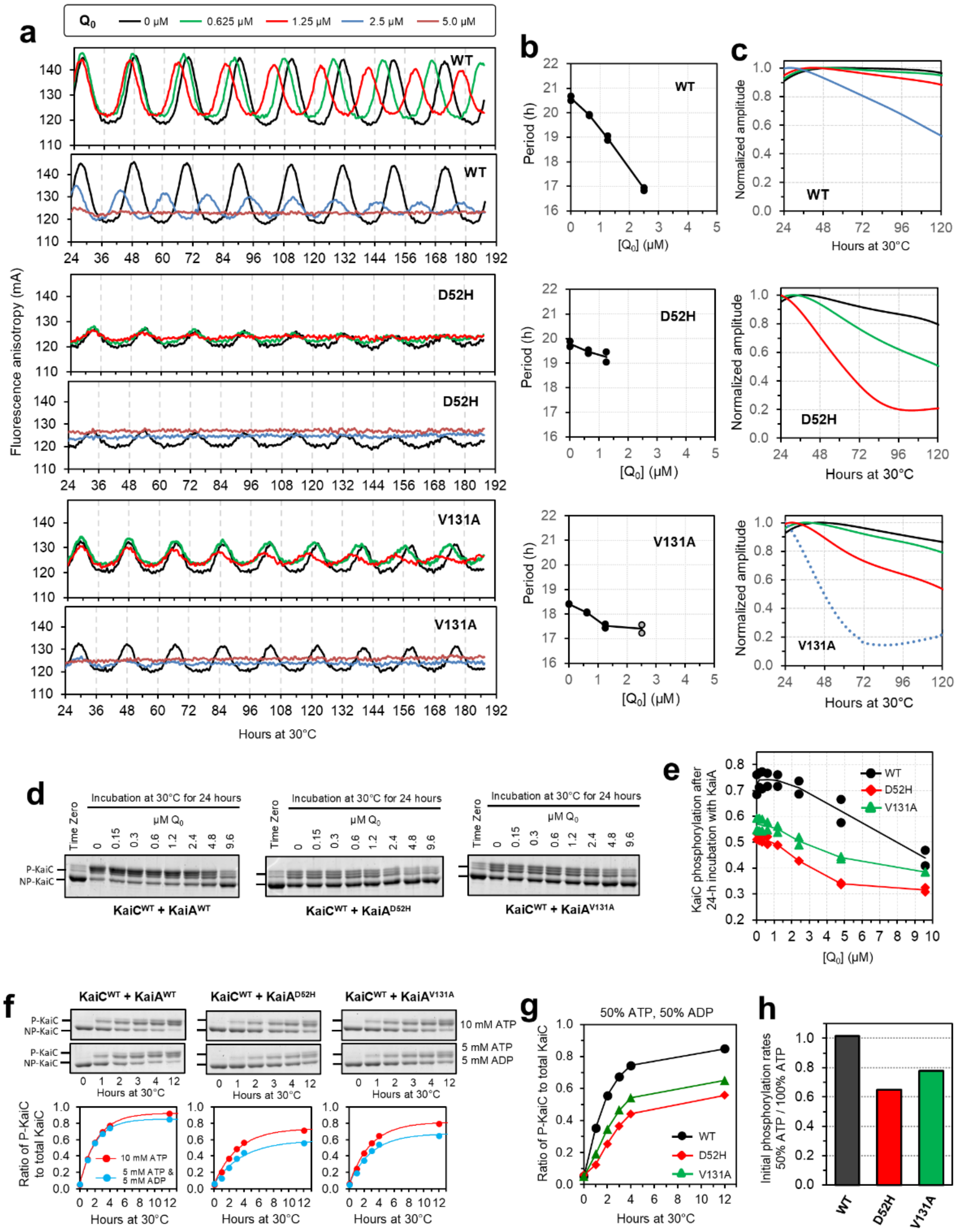

**Extended Data Fig. 5. D52H and V131A mutations of KaiA have significant impact upon biochemical properties. a.** *In vitro* oscillations reconstituted with KaiC, KaiB, and the wild-type (KaiA[WT]) or mutant variants (D52H or V131A) of KaiA with various concentrations (0, 0.625, 1.25, 2.5, 5 μM) of $Q_0$. The oscillations were monitored by fluorescence anisotropy, which measures the change in rotational diffusion of fluorescein-labelled KaiB as a monitor of the association/dissociation of the KaiABC complex.[58] Reactions without $Q_0$ were treated with 0.05 % ethanol as a vehicle control. (n = 2). **b.** Estimated period length of the *in vitro* KaiABC oscillator (IVO/PTO) in the presence of $Q_0$. **c.** Damping of the amplitude of the *in vitro* oscillations over time, measured from the instantaneous amplitude (normalized to the maximum amplitude value after 24 h). **d.** Representative gel images of KaiC incubated with KaiA and various concentrations of $Q_0$. Dephosphorylated KaiC (3.5 μM) was incubated with 1.5 μM of $KaiA^{WT}$, $KaiA^{D52H,}$ or $KaiA^{V131A}$ and with 0-9.6 μM $Q_0$ at 30°C for 24 hours, and the phosphorylation state of KaiC was analyzed by SDS-PAGE (lowest band is unphosphorylated KaiC, upper bands are various phosphorylation states of KaiC. **e.** KaiC phosphorylation levels after 24 hours of incubation with KaiA (unnormalized data plots corresponding to Fig. 3e, but here including 10 μM $Q_0$). Data from two independent experiments are plotted. **f.** D52H and V131A mutations in *kaiA* result in lower activity of KaiA protein to stimulate KaiC auto-phosphorylation, and alter KaiA's sensitivity to ATP/ADP ratio. Representative gel images of the time-course of KaiC phosphorylation with $KaiA^{WT}$, $KaiA^{D52H}$ or $KaiA^{V131A}$. KaiC protein was incubated with $KaiA^{WT}$, $KaiA^{D52H}$, or $KaiA^{V131A}$ proteins either in the presence of 10 mM ATP (100% ATP) or 5 mM ATP/5 mM ADP (50% ATP). **g.** Time-course of KaiC auto-phosphorylation in the presence of the wild-type or variants of KaiA with 5 mM ATP and 5 mM ADP (50% ATP). **h.** Effect of ATP/ADP ratio on KaiA-stimulated KaiC autophosphorylation in the presence of the WT or TC-variants of KaiA.

**Extended Data Fig. 6**

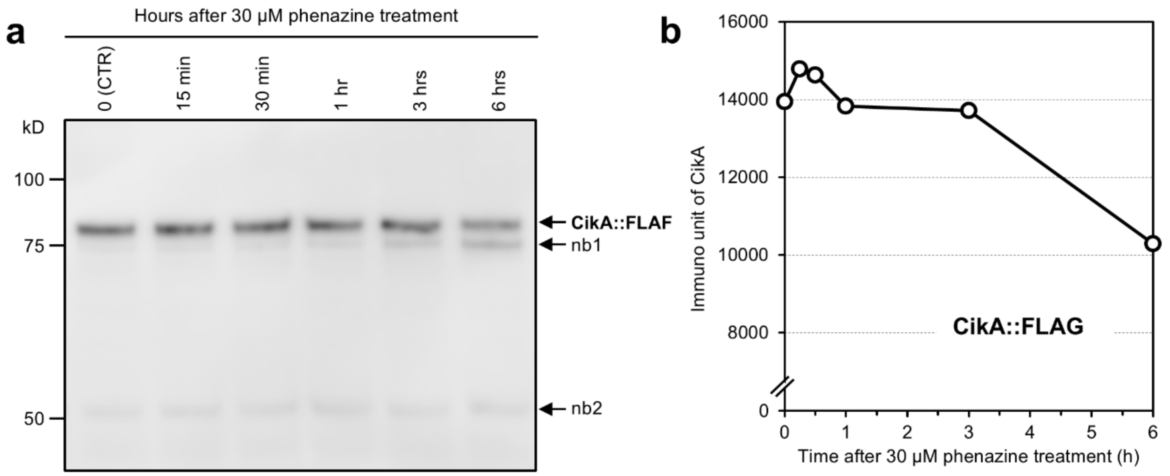


**Extended Data Fig. 6. CikA levels are sensitive to oxidizing conditions.** To evaluate the sensitivity of CikA protein to redox changes in living cells with an immunoblot assay, FLAG-tagged CikA strain was generated with FLAG-tagged CikA inserted in the neutral site 4 (NS4) of a ΔCikA strain. The FLAG-tagged CikA fully restored WT properties to the ΔCikA strain. FLAG-tagged CikA levels decline after cyanobacterial cells are treated with 30 μM phenazine. **a.** After treatment of the CikA::FLAG-expressing cells with 30 μM phenazine, extracts were prepared from the cells and proteins separated on 8% SDS-PAGE, followed by immunoblot assay with FLAG antibody. The CikA band is indicated as “CikA::FLAG,” and non-specific bands are labeled as “nb1” and “nb2.” **b.** Densitometric quantification of the CikA::FLAG band on the immunoblot depicted in panel **a**.

**Extended Data Fig. 7**

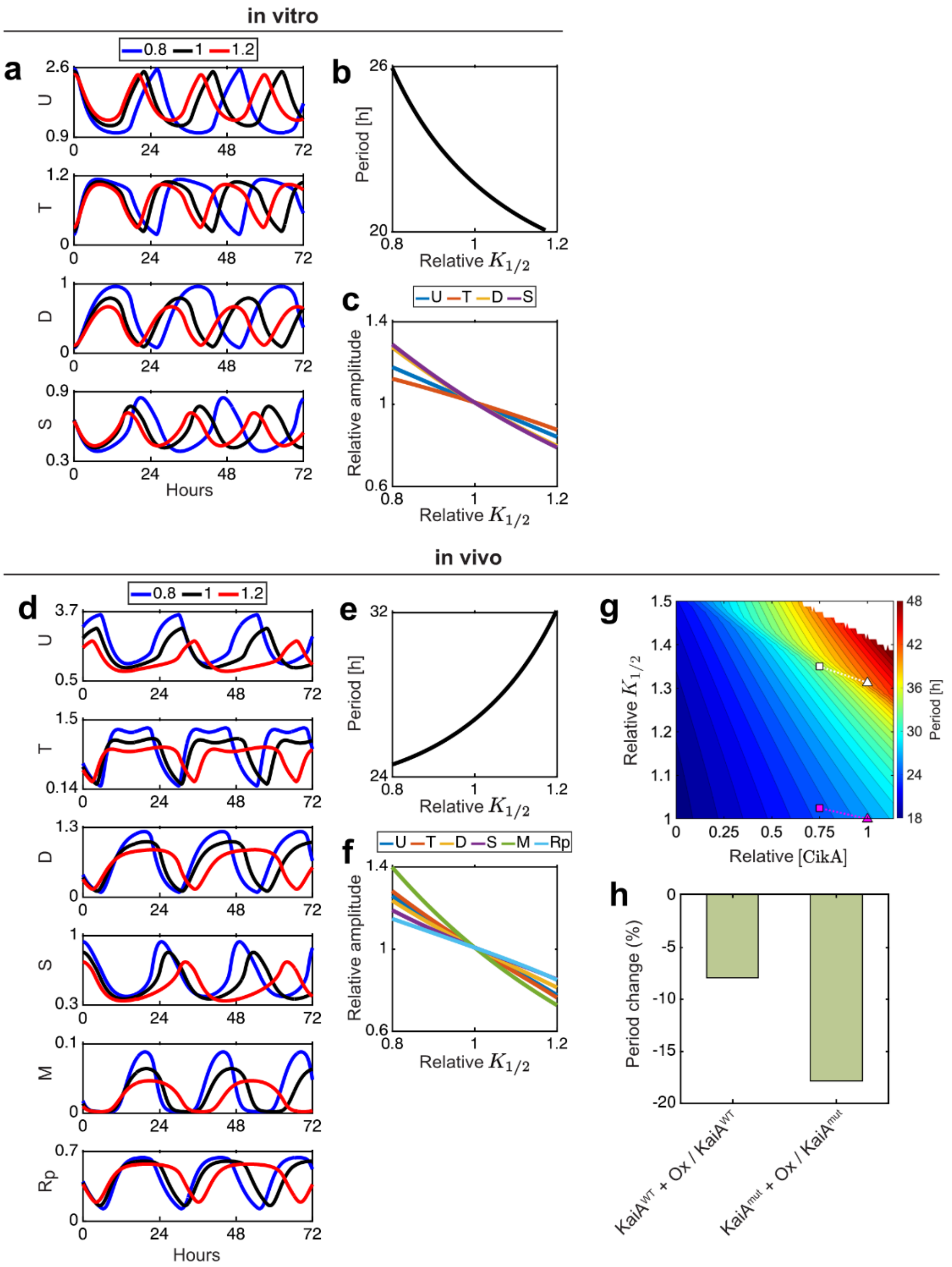


**Extended Data Fig. 7. Simulation of *in vivo* and *in vitro* KaiC phosphorylation cycles and *in vivo* rhythms of transcription in wild-type vs. KaiA-mutant strains under standard conditions and under oxidizing conditions. a** and **d.** Time series of variables in the mathematical models for the KaiC phosphorylation cycle under (**a**) *in vitro* and (**d**) *in vivo* conditions with different values of $K_{1/2}$ for KaiA activity ($K_{1/2}$ = 0.8 {blue}, 1.0 {black}, 1.2 {red}). Panels from top to bottom: unphosphorylated KaiC (U),

KaiC phosphorylated at T432 (T), KaiC phosphorylated at both T432 and S431 (D, aka ST), and KaiC phosphorylated at S431 (S). **d.** In addition to the time series of the KaiC phosphorylation cycle shown in **a**, the time series of transcription (as *kaiC* mRNA {M}), and phosphorylated RpaA (Rp) levels are also shown. **b** and **e**. Dependence of the period of the oscillations on relative $K_{1/2}$ of KaiA activity *in vitro* (**b**) and *in vivo* (**e**). Increasing the value of $K_{1/2}$ shortens the period *in vitro*, whereas it lengthens the period *in vivo*. This occurs because the amplitude of the IVO/PTO oscillator is dramatically reduced and its period is slightly shortened *in vitro* with $KaiA^{D52H}$ and $KaiA^{V131A}$ proteins because their activity on KaiC autophosphorylation is reduced. However, the periods of the simulated PTO/TTFL oscillations are lengthened with the $KaiA^{D52H}$ and $KaiA^{V131A}$ proteins because the lower amplitude of the embedded PTO leads to a lower concentration of KaiC in its D form, thereby leading to a slower/lower activation of SasA, and therefore a slower activation of RpaA phosphorylation. **c** and **f**. Dependence of the amplitude of each variable on relative $K_{1/2}$ of KaiA activity *in vitro* (**c**) and *in vivo* (**f**). The colors of the lines denote the different variables as indicated in the keys above panels (**c**) and (**f**). We set $K_{1/2} = 0.43$ as the $KaiA^{WT}$ value *in vitro* in (**b**) and (**c**), and $K_{1/2} = 0.344$ as the $KaiA^{WT}$ value *in vivo* in (**e**) and (**f**). The horizontal axis is normalized by these wildtype ($KaiA^{WT}$) values. The amplitudes in (**c**) and (**f**) are also normalized by the wildtype amplitudes. **g.** Period of the *in vivo* KaiC oscillation as a function of (i) relative $K_{1/2}$ of KaiA and (ii) level of CikA [CikA] values. Magenta and white triangles indicate the $K_{1/2}$/CikA values for the $KaiA^{WT}$ (magenta) and KaiA mutants (white) under the standard (non-oxidizing) conditions depicted in Fig. 3d. Magenta and white squares are the values for the $KaiA^{WT}$ (magenta) and KaiA mutants (white) under oxidizing conditions. The horizontal axis is normalized by the CikA levels under standard (non-oxidizing) conditions. The vertical axis is normalized by the value of $K_{1/2}$ for $KaiA^{WT}$ under standard (non-oxidizing) conditions. Contour lines indicate one-hour period differences. Oxidizing conditions are simulated as a reduction of CikA levels, based on the experimental data of Extended Data Fig. 6 and the data reported by Ivleva *et al.*[36] **h**. Percent period changes in wildtype ($KaiA^{WT}$) vs. KaiA mutant strains elicited by oxidization (simulated by reduction in [CikA]). Also see Figure 3D for rhythm traces based upon this simulation. We used parameter values indicated by the symbols (squares and triangles) in (**g**). The values of all the parameters for simulations are listed in Extended Data Tables 1 and 2.

#### Extended Data Fig. 8

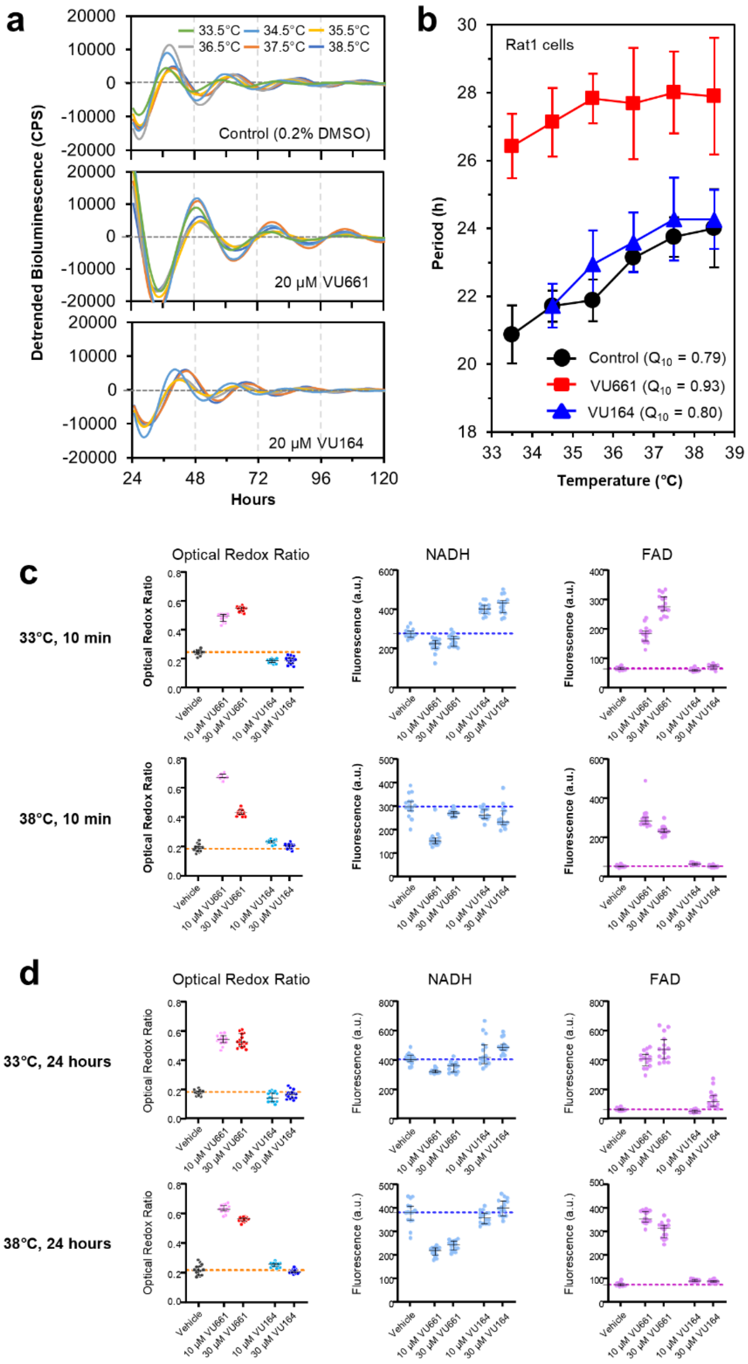


**Extended Data Fig. 8. Disrupting the redox balance in mammalian cells impairs temperature compensation. a.** Effect of phenazine carboxamide VU661 on circadian luminescence rhythms (Bmal1-Luc reporter) at different temperatures in mammalian Rat-1 cells.[38] Dexamethasone-synchronized Rat-1 Bmal1-Luc cells were exposed to 20 μM of the phenazine carboxamide compound VU661 or its inactive analog (isomeric benzquinoline carboxamide VU164), both of which were dissolved in DMSO (final concentration of DMSO was 0.2%). The luminescence rhythms were measured at various temperatures as described in the Methods. Cells treated with 0.2% DMSO were used as the control. **b.** Mean period lengths of circadian luminescence rhythms are plotted versus temperature, with bars representing means ±S.D. from three biologically independent experiments. $Q_{10}$ of Rat-1 cells over a temperature gradient of 33-39°C with and without VU661/VU164 treatment are calculated. Note that while the $Q_{10}$ of U2OS cells is > 1.0 (Fig. 5b), the $Q_{10}$ of Rat-1 cells is < 1.0, but in both cases treatment with VU661 alters $Q_{10}$ values. **c.** ORR, NADH, and FAD values for all the analyzed U2OS cell images after the 10 min treatment at 33°C and 38°C. Each dot in the graph is the value extracted from one image containing 10-40 cells, n = 15 for each group. The black bars indicate the median value of each group, and the error bar represent the interquartile range of each group. The dashed line indicates the median value for the control group (vehicle). **b.** ORR, NADH, and FAD values for all the analyzed U2OS cell images after the 24 h treatment at 33°C and 38°C. These ORR data are replotted in Fig. 4b. Each dot in the graph is the value extracted from one image containing 10-40 cells, n = 15 for each group. The black bars indicate the median value of each group, and the error bar represent the interquartile range of each group. The dashed line indicates the median value for the control group (vehicle).

**Extended Data Fig. 9**

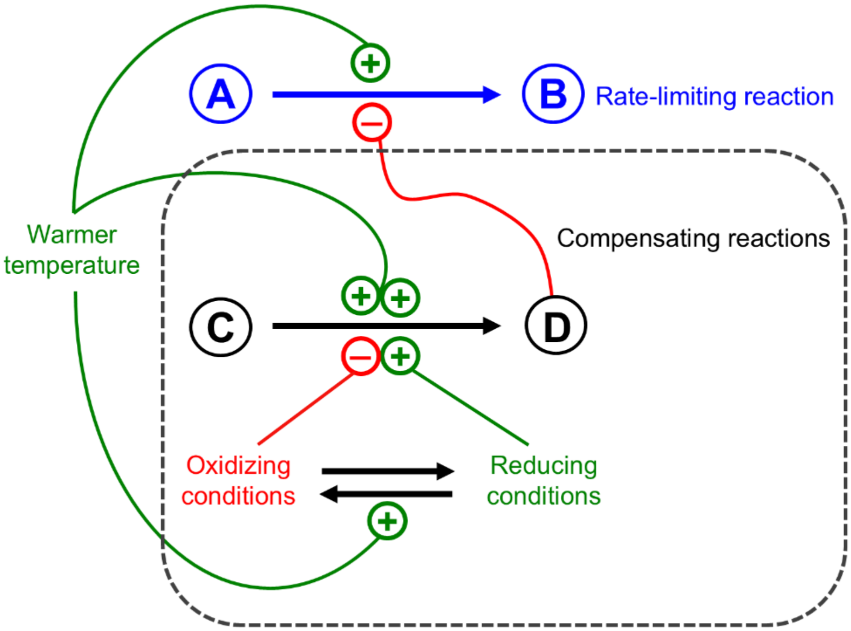


**Extended Data Fig. 9. New model of temperature compensation based upon redox sensing as an integral component of temperature compensation.** The "Compensation Reaction" of the model in Figure S1A is revealed to be regulated by both redox and temperature. Warmer temperature strongly enhances the rate of the C $\rightarrow$ D compensation reaction, but it also oxidizes the cellular milieu, which serves to slow the C $\rightarrow$ D reaction, ultimately balancing the rate-limiting A $\rightarrow$ B reaction to maintain a constant circadian period. Metabolic conditions that could modify period also affect redox, and this perturbation is corrected via redox sensing/signaling to the A $\rightarrow$ B reaction. The redox sensors in cyanobacteria include KaiA and CikA, and the KaiA TC mutants identified here are hyper-sensitive to redox changes so that the counterbalancing mechanism is inaccurate. Consequently, in cells expressing the mutant KaiA proteins, oxidizing conditions impair the TC mechanism so that circadian period is no longer conserved across a physiological range of temperatures.

**Extended Data Fig. 10**

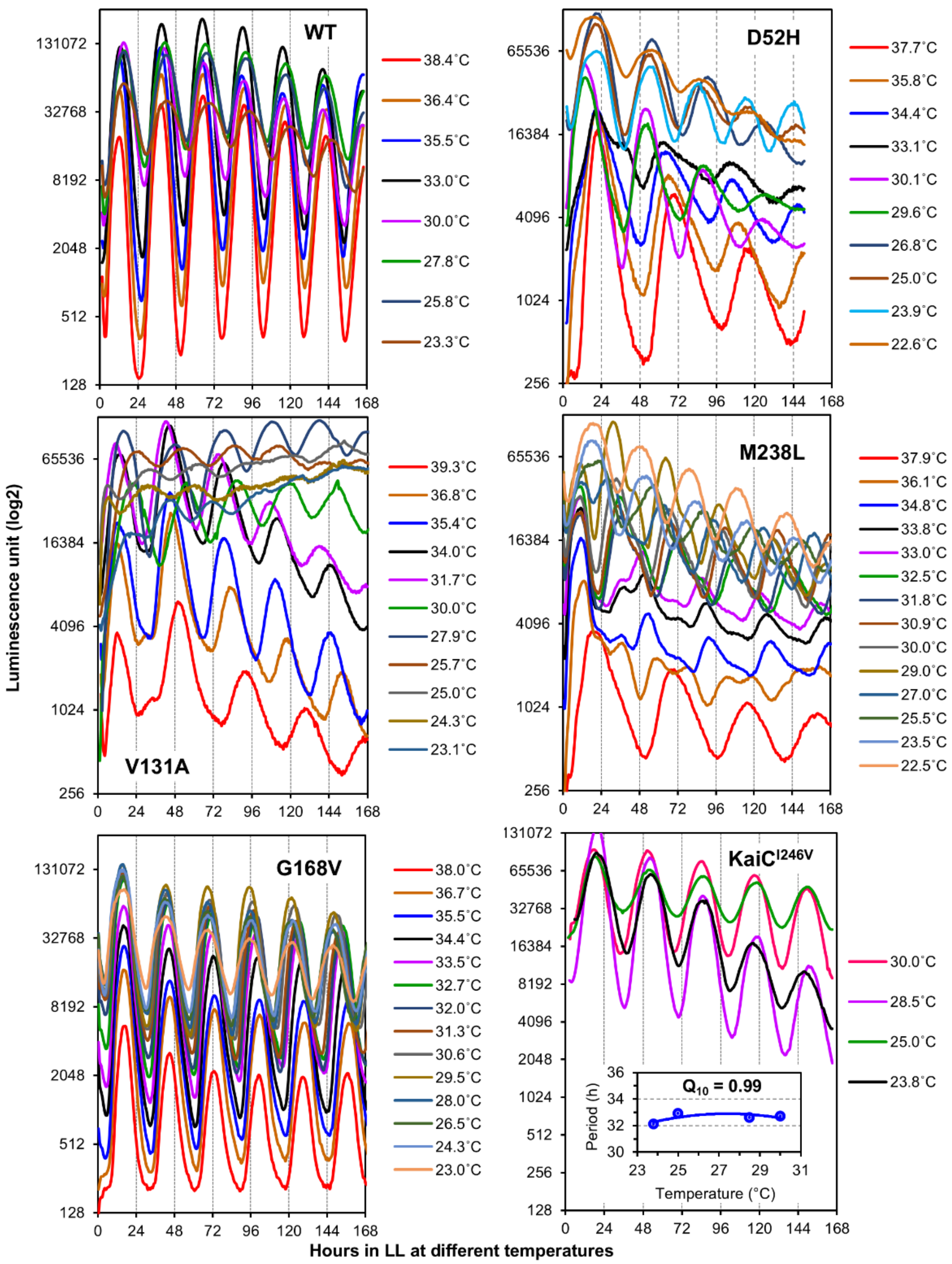


**Extended Data Fig. 10. Representative raw-data circadian luminescence rhythm traces for key strains over a range of temperatures.** Luminescence rhythms are shown for wild type, *kaiA* TC mutants (D52H, V131A, M238L, and G168V), and the long period KaiC mutant I246V in LL at different temperatures. As is true for wild-type, the long-period $KaiC^{I246V}$ mutant is temperature compensated ($Q_{10}$ = 0.99 for 23°C to 30°C).

### Supplementary Information

**Supplementary Fig. 1**

**Supplementary Tables 1 and 2**

#### Supplementary Fig. 1

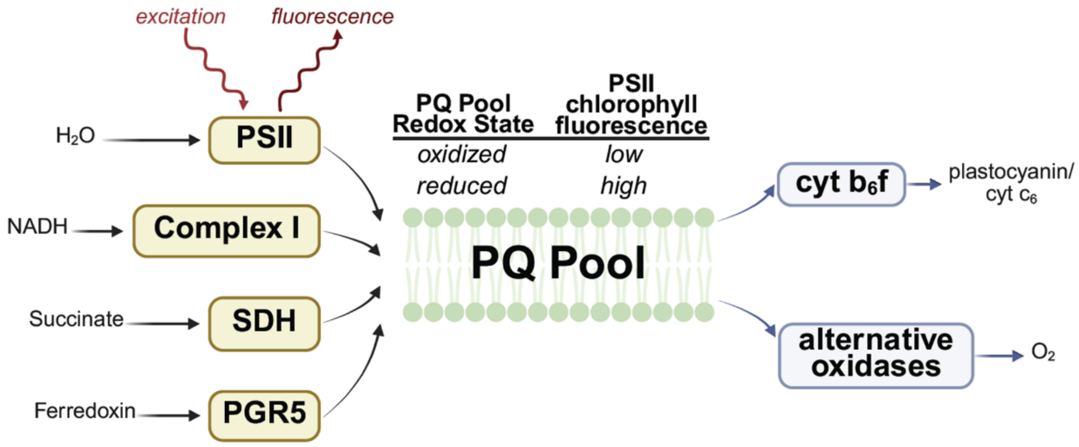


**Supplementary Fig. 1. Photosynthesis and respiration share a common plastoquinone (PQ) pool in cyanobacteria.** The pool is reduced (black arrows) by Photosystem II (PSII), Photosynthetic Complex I, Succinate Dehydrogenase (SDH), and Proton Gradient Regulator 5 (PGR5). The pool is oxidized (blue arrows) by cytochrome (cyt) $b_6f$ and various alternative oxidases such as cyt bd and PQ terminal oxidase (PTOX). The electron source or sink for each of these complexes is soluble, thus connecting the cytosolic and membrane redox states. PSII contains chlorophyll molecules that are excited by visible light and may relax via fluorescence. When the PQ pool is in a reduced state, the yield of PSII chlorophyll fluorescence is high. When the PQ pool is in an oxidized state, the yield of PSII chlorophyll fluorescence is low. This variable chlorophyll fluorescence signal provides a nonintrusive method to monitor intracellular redox in cyanobacteria. This figure was created in BioRender.

**Supplementary Table 1. List of parameter values for numerical simulations**

| Description | References | Parameter name | Value | Unit |
|---|---|---|---|---|
| $U \rightarrow T$ (Basal rates) | Rust *et al.* 2007 | $k_{UT}^{0}$ | 0 | $h^{-1}$ |
| $T \rightarrow D$ (Basal rates) | Rust *et al.* 2007 | $k_{TD}^{0}$ | 0 | $h^{-1}$ |
| $S \rightarrow D$ (Basal rates) | Rust *et al.* 2007 | $k_{SD}^{0}$ | 0 | $h^{-1}$ |
| $U \rightarrow S$ (Basal rates) | Rust *et al.* 2007 | $k_{US}^{0}$ | 0 | $h^{-1}$ |
| $T \rightarrow U$ (Basal rates) | Rust *et al.* 2007 | $k_{TU}^{0}$ | 0.21 | $h^{-1}$ |
| $D \rightarrow T$ (Basal rates) | Rust *et al.* 2007 | $k_{DT}^{0}$ | 0 | $h^{-1}$ |
| $D \rightarrow S$ (Basal rates) | Rust *et al.* 2007 | $k_{DS}^{0}$ | 0.31 | $h^{-1}$ |
| $S \rightarrow U$ (Basal rates) | Rust *et al.* 2007 | $k_{SU}^{0}$ | 0.11 | $h^{-1}$ |
| $U \rightarrow T$ (Maximal effect of KaiA) | Rust *et al.* 2007 | $k_{UT}^{A}$ | 0.479 | $h^{-1}$ |
| $T \rightarrow D$ (Maximal effect of KaiA) | Rust *et al.* 2007 | $k_{TD}^{A}$ | 0.213 | $h^{-1}$ |
| $S \rightarrow D$ (Maximal effect of KaiA) | Rust *et al.* 2007 | $k_{SD}^{A}$ | 0.5057 | $h^{-1}$ |
| $U \rightarrow S$ (Maximal effect of KaiA) | Rust *et al.* 2007 | $k_{US}^{A}$ | 0.0532 | $h^{-1}$ |
| $T \rightarrow U$ (Maximal effect of KaiA) | Rust *et al.* 2007 | $k_{TU}^{A}$ | 0.0798 | $h^{-1}$ |
| $D \rightarrow T$ (Maximal effect of KaiA) | Rust *et al.* 2007 | $k_{DT}^{A}$ | 0.173 | $h^{-1}$ |
| $D \rightarrow S$ (Maximal effect of KaiA) | Rust *et al.* 2007 | $k_{DS}^{A}$ | -0.32 | $h^{-1}$ |
| $S \rightarrow U$ (Maximal effect of KaiA) | Rust *et al.* 2007 | $k_{SU}^{A}$ | -0.133 | $h^{-1}$ |
| Concentration of KaiA causing half-maximal effect on KaiC (*in vitro* wild type) | Rust *et al.* 2007 | $K_{1/2}$ | 0.43 | μM |
| Concentration of KaiA causing half-maximal effect on KaiC (*in vivo* wild type) | This study | $K_{1/2}$ | 0.344 | μM |
| Concentration of KaiA | Rust *et al.* 2007 | $A_T$ | 1.3 | μM |
| Total concentration of KaiC *in vitro* | Rust *et al.* 2007 | $[\mathrm{KaiC}]$ | 3.4 | μM |
| Degradation rate of KaiC proteins | Modified from Teng *et al.* 2013 | $V_d$ | 0.021 | $h^{-1}$ |
| Translation rate of KaiC protein | Modified from Teng *et al.* 2013 | $K_s$ | 2.8 | $h^{-1}$ |
| Maximum transcription rate of *kaiC* mRNA | Modified from Teng *et al.* 2013 | $V_s$ | 1.785 | $\mu M \cdot h^{-1}$ |
| Threshold constant determining the RpaA levels for the half maximum value of transcription | Modified from Teng *et al.* 2013 | $K_i$ | 1 | μM |
| the Hill coefficient | Modified from Teng *et al.* 2013 | $n$ | 6 | - |
| Degradation rate of *kaiC* mRNA | Modified from Teng *et al.* 2013 | $V_m$ | 1.2 | $h^{-1}$ |
| Total RpaA levels | This study | $R_T$ | 1 | μM |
| Phosphorylation rate of RpaA | This study | $k_1$ | 0.6 | $\mu M^{-1} \cdot h^{-1}$ |
| Basal dephosphorylation rate of RpaA | This study | $k_2$ | 0.3 | $h^{-1}$ |
| Dephosphorylation rate of RpaA | This study | $k_3$ | 0.21 | $\mu M^{-2} \cdot h^{-1}$ |
| CikA levels (non-oxidized condition) | This study | $[\mathrm{CikA}]$ | 1.32 | μM |

**Supplementary Table 2. Values of parameters to model the KaiA mutant and oxidizing conditions**

| Conditions | $K_{1/2}$ | $[\mathrm{CikA}]$ |
|---|---|---|
| *in vitro* wildtype, $KaiA^{WT}$ | 0.43 | - |
| *in vitro* KaiA mutant $KaiA^{mut}$ | 0.6063 | - |
| *in vivo* wildtype non-oxidized, $KaiA^{WT}$ | 0.344 | 1.32 |
| *in vivo* wildtype oxidized, $KaiA^{WT}$ + Ox | 0.3526 | 0.99 |
| *in vivo* KaiA mutant non-oxidized, $KaiA^{mut}$ | 0.4515 | 1.32 |
| *in vivo* KaiA mutant oxidized, $KaiA^{mut}$ + Ox | 0.4644 | 0.99 |
